# How Intrinsic Neural Timescales Relate To Event-Related Activity – Key Role For Intracolumnar Connections

**DOI:** 10.1101/2025.01.10.632350

**Authors:** Yasir Çatal, Angelika Wolman, Andrea Buccellato, Kaan Keskin, Georg Northoff

**Author notes:** These authors contributed equally (Angelika Wolman, Andrea Buccellato, Kaan Keskin). Correspondence: Yasir Çatal.

## Abstract

The relationship of the brain’s intrinsic neural timescales (INTs) during the resting state with event-related activity in response to external stimuli remains poorly understood. Here, we bridge this gap by combining computational modeling with magnetoencephalography (MEG) data to investigate the relation of intrinsic neuronal timescales (INT) with task-related activity, e.g., event-related fields (ERFs). Using the Jansen-Rit model, we first show that intracolumnar (and thus intra-regional) excitatory and inhibitory connections (rather than inter-regional feedback, feedforward and lateral connections between the columns of different regions) strongly influence both resting state INTs and task-related ERFs. Secondly, our results demonstrate a positive relationship between the magnitude of event-related fields (mERFs) and INTs, observed in both model simulations and empirical MEG data collected during an emotional face recognition task. Thirdly, modeling shows that the positive relationship of mERF and INT depends on intracolumnar connections through observing that the correlation between them disappears for fixed values of intracolumnar connections. Together, these findings highlight the importance of intracolumnar connections as a shared biological mechanism underlying both the resting-state’s INTs and the task-state’s event-related activity including their interplay.

## Introduction

The brain operates as a dynamic system, constantly interacting with and adapting to the environment. While responding to external stimuli is crucial, the brain also generates spontaneous activity in the absence of explicit tasks or goals. This intrinsic activity is characterized by intrinsic neural timescales (INTs), which describe the temporal dynamics of neural processes^1^. INTs have been extensively studied in both resting-state and task-related contexts^2–5^. Despite this, a critical gap remains: how do INTs, as markers of intrinsic brain activity and its timescales, relate to event-related activity elicited by external stimuli? In this paper, we bridge this gap using computational modeling and magnetoencephalography (MEG) data.

The heterogeneity of INTs across the brain is well-documented^6–8^. INTs represent the time window during which prior information influences the processing of new stimuli, enabling temporal integration^2,5,7^. Neuroimaging studies have linked INTs to diverse cognitive functions, such as reward processing^8^, self-referential thought^9,10^, narrative comprehension^11^, and consciousness^12–14^. Furthermore, it is widely recognized that various task paradigms and types of stimulation induce evoked neural activity, such as event-related potentials or fields. ^15–17^). In contrast, how the resting state INTs relate to the task-state evoked activity is an important question that so far remains unanswered.

Our focus on the relationship between INTs and event-related fields (ERFs) is motivated by three complementary lines of reasoning. First, both INTs and ERFs are independently linked to neural computations underlying behavior^18,19^. Despite this, the interplay between them remains largely unexplored. Second, prior research demonstrates non-additive interactions between resting-state/prestimulus activity and task-evoked/poststimulus activity, observed across functional MRI (fMRI)^20–32^, EEG^4,9,33–39^, and single-unit recordings^40–43^. The findings of both non-additive rest-task interaction and the relationship of behavior with both ERF and INT suggest that intrinsic activity and its INT may be related to task-related neural responses, the underlying mechanisms of their relation which remains unclear. Finally, on the theoretical side, the fluctuation-dissipation theorem^44–46^ offers a theoretical framework for their interaction, positing that the temporal structure of a system’s spontaneous activity (specifically, its autocorrelation function which INT is usually calculated from) predicts its average response to external inputs (e.g. ERF). Together, these points strongly support the assumption of an interplay between INTs and ERFs.

The goal of this study is to uncover the relationship between the resting state’s intrinsic neural timescales (INTs) and event-related activity. To operationalize these concepts, we define INTs by using the lag at which the autocorrelation function (ACF) reaches half of its original value (ACW-50). While event-related activity is determined as the magnitude of event-related fields (mERFs) — stimulus-locked neural activity averaged across trials. This relationship will be examined through a combination of computational modeling and empirical data analysis. We first address our research question using the computational Jansen-Rit model^47–51^, which simulates the dynamics of coupled cortical columns. The model provides a mechanistic link to empirical measures by approximating the average depolarization of pyramidal neurons within cortical columns, the primary source of EEG/MEG signals^47,48,50^ which we here use to measure both INT and ERF including their relationship on empirical grounds.

Our initial aim is to explore how INTs and mERFs depend on structural connections such as intra-columnar/intra-regional connections and inter-columnar/inter-regional connections (i.e. lateral, feedback and feedforward connections^50^) within and between cortical columns. Specifically, we sweep a range of values for each model parameter while keeping others constant and evaluate the relationship between the different intra-and inter-columnar structural connections and mERF/INTs. By identifying parameters that simultaneously influence both INTs and mERFs, we aim to provide a shared biological basis for both resting state INTs and task-related activity/ERF as suggested by their close relationship on both neuronal and behavioral levels.

Recent evidence on large-scale cortical gradients suggests that intraregional (intracolumnar) excitatory and inhibitory connections correlate positively with intrinsic neural timescales^52^; this motivates our hypothesis that intracolumnar connections (as distinguished from feedback, feedforward and lateral connections) will have the greatest impact on both INTs and mERFs. Thus, our first aim gives us a structural parameter to investigate the possibly shared computational basis of mERF and INT.

Next, our second aim focuses on the relationship of INTs and mERFs in the computational model. The background rationale is as follows. If INTs and mERFs share a computational basis in their underlying structural parameter (like intracolumnar connections), one expects to see a correlation between mERFs and INTs; this is further reinforced on the theoretical side by the fluctuation-dissipation theorem which states that the linear response of a system to an external input is proportional to its autocorrelation function^44–46^ which provides the basis for our INT estimation.

The third aim of the paper is testing the insights obtained from modeling in empirical data. We use an open-access MEG data set providing both resting state and a task state, that is, the Hariri-Hammer task for emotional face recognition MEG data^53^. We estimate INTs in resting-state MEG data using the ACW metric, consistent with the computational model; while, for task-related data, we calculate the mERF. Based on previous findings^54–56^, we anticipate that significant electrophysiological activity measured by the mERF will emerge primarily in temporal MEG sensors and within the 100–200 ms window after trial onset. These sensors will then be used to calculate ACW. Subsequently, the values obtained from these calculations will be correlated with the previously derived mERFs, to connect the empirical findings with the computational observations.

Our findings are as follows: (i) computational: intracolumnar excitatory and inhibitory connections (rather than feedback, feedforward and lateral connections between the columns of different regions) are the only parameter that correlates with both INTs and mERFs, (ii) computational: increased intracolumnar connections lead to both longer ACW and stronger mERF responses, supporting the hypothesized link between INTs and mERFs; (iii) computational: positive relationship between INTs and mERFs depend on the degree of the underlying intracolumnar excitatory and inhibitory connections; (iv) empirical: MEG data reveal the most significant contrasts between emotional face and non-emotional shape conditions in temporal channels during both encoding and probe conditions, specifically at 100 ms and 250 ms after stimulus onset; (v) empirical: INTs and mERFs are correlated positively in the MEG data, consistent with the modeling results. These findings are summarized in Figure 1, which provides an overview of our workflow and key results. Together, our work systematically integrates computational and empirical approaches to uncover how the brain’s intracolumnar excitatory and inhibitory connections mediate the relationship between the resting state’s intrinsic neural timescales and event-related activity during external stimulation.

**Figure 1.**
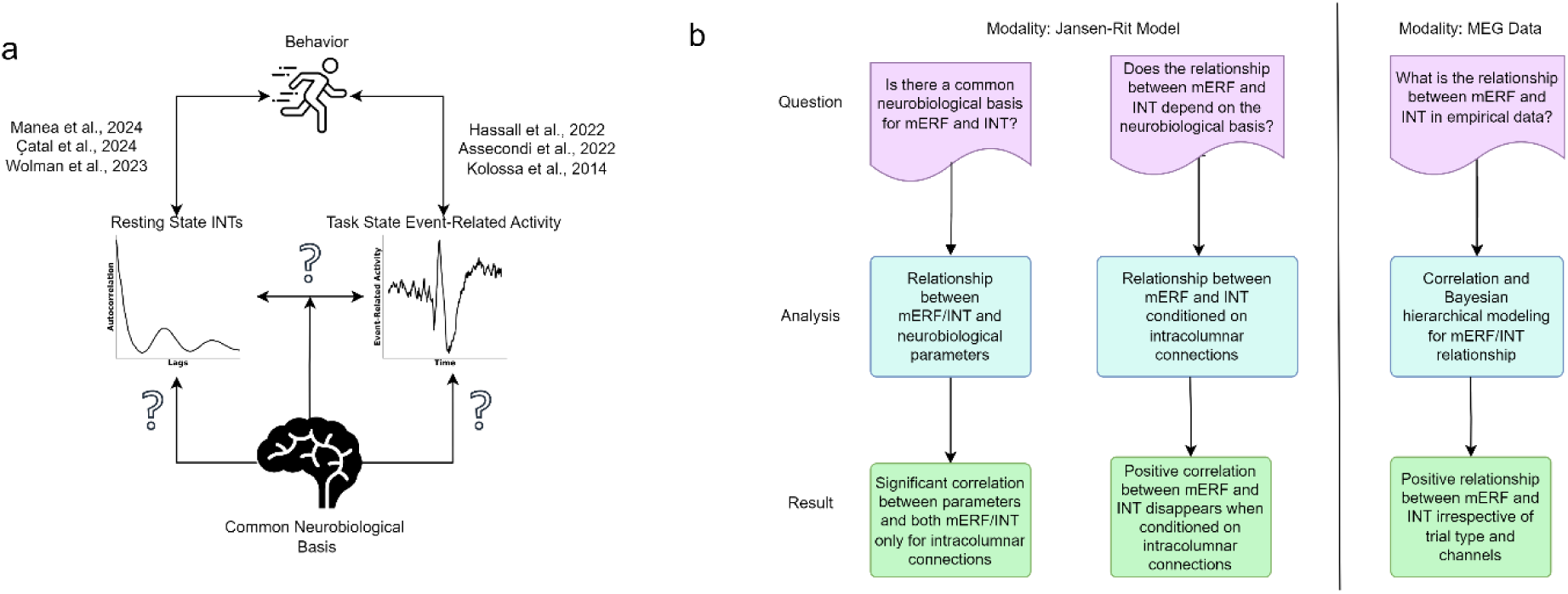
Outline of the research. a. Previous research indicate both resting state INTs and event-related activity at task state is related to behavior. Our research focuses on the relationship between INTs and event-related activity and a possible common neurobiological basis between them. b. The questions, analyses and main results of our research.

## Methods

### Jansen-Rit Model of Coupled Cortical Columns

Jansen-Rit model is a mathematical model of ERP/F generation. The model follows a basic microcircuit observed across species^57^. In this microcircuit, the pyramidal cells receive excitatory input from extrinsic afferent systems and spiny cells; and inhibitory input from GABAergic interneurons. A cortical area in the model is described by these three populations: pyramidal cells, excitatory interneurons and inhibitory interneurons. Pyramidal cells receive input from inhibitory and excitatory interneurons in the same population and send output to inhibitory and excitatory interneurons. Excitatory interneurons can be thought as spiny stellate cells in layer 4 and receive feedforward connections^58^ from external cortical areas. Excitatory pyramidal cells and inhibitory interneurons can be found in supragranular and infragranular layers respectively and receive input from backward and lateral connections formed with external cortical areas. We used two cortical columns (as representing two distinct regions) for the results presented in figure 3. The general model equations in hierarchical form are described as follows:

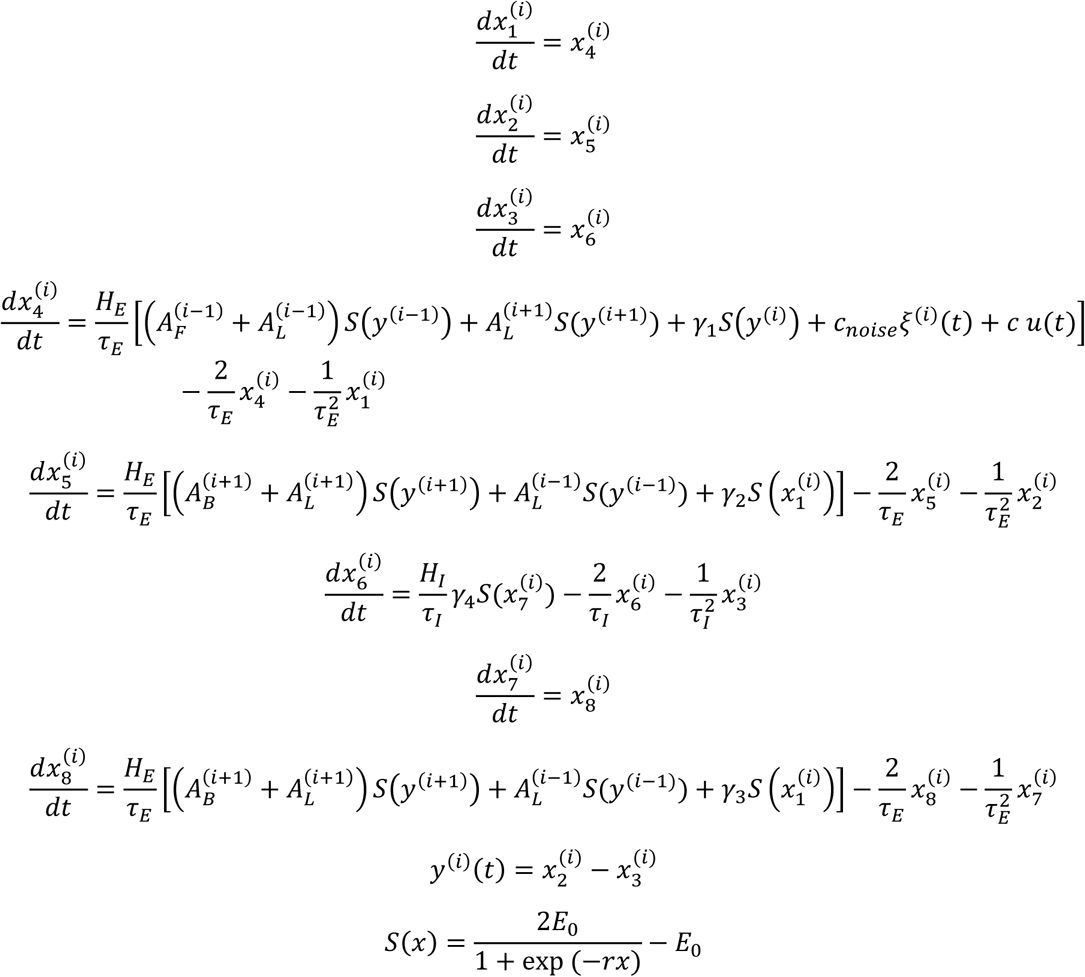

Where *i* denotes the specific column in the hierarchy and goes from 1 to *n* where *n* denotes the number of regions in the hierarchy. Hence by definition *a*^(0)^ = *a*^(*n*+1)^ = 0 where *a* is any variable in the model. The variable *y* corresponds to the picked-up signal by EEG / MEG and is the average depolarization of pyramidal cells. The parameters *A*_*B*_, *A*_*F*_ and *A*_*L*_ corresponds to backward, forward and lateral connections between coupled cortical columns. *ξ* is gaussian white noise, corresponding to background activity, with variance *c*_*noise*_= 0.1. *u*(*t*) is external input, a delta function fired at trial onsets. *c* scales the strength of this delta function and is set to 10^4^. The parameters *H*_*E*,*I*_ tune the maximum amplitude of post – synaptic potentials. *τ*_*E*,*I*_ represent the rate constants of cell membrane. Following David and Friston^50^, values for *H* and *τ* are set to *H*_*E*_=3.25, *H*_*I*_=29.3, *τ*_*E*_=10 ms, *τ*_*I*_=15 ms. *γ* parameters scale the intracolumnar connections. Empirical observations constrain the values of *γ* with relations *γ*_2_ = 0.8*γ*_1_, *γ*_3_ = *γ*_4_ = 0.25*γ*_1_. In our study, the default value of *γ*_1_ was set to 50 and *A*_*F*_ was set to 2.5 to ensure even when all other *A*_*i*_ is 0, the input can still be transferred between the two areas. The function *S*(*x*) transforms the average membrane potential of a subpopulation into an average firing rate. It is parametrized by *E*_0_ and *r* which were set to 2.5 and 0.56 respectively.

The stochastic differential equations were solved using DifferentialEquations.jl package in Julia using adaptive step-size SKenCarp solver^59–62^. In the resting state, we simulate 100 seconds of data, then calculate one autocorrelation window (ACW) value from each of the 10 seconds of data, and finally average the ACW values to get one ACW per region. In the task state, we give a delta-function input every 5 seconds as in the empirical task. We gave the input 19 times, first at second 5 and last at second 95, each corresponding to a trial. ERF was calculated as the average of these time – locked trials. The magnitude of ERF was calculated as the root-mean squared values of ERF averaged over 0.7 seconds after the trial onset.

### Estimation of Intrinsic Neural Timescales

Autocorrelation function is used to estimate intrinsic neural timescales (INT) from both the simulations and neural recordings. The autocorrelation function (ACF) *r*(*l*) of a signal x is the signal’s correlation with itself on different lags:

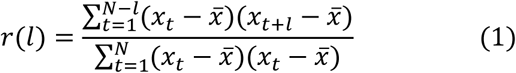

where *l* denotes lag, *x̄* denotes the mean of *x* and *N* is the number of sampling points. In practice, the ACF is calculated as the inverse Fourier transform of power spectrum, utilizing Wiener-Khinchine theorem for computational efficiency^63^ when there is no missing data. We used autocorrelation window (ACW) to get one value representative of INT from the ACF. ACW is defined as the first lag below *r* = 0.5.

The ACW values were estimated using a sliding window approach. From every non-overlapping 10 second segment of the time-series, we calculated one ACW value, then averaged them to get a more reliable estimate of INTs in both simulations and empirical data.

### Data Acquisition, Task Design and Preprocessing

NIMH intramural healthy volunteer dataset^53^ is a large-scale open-access dataset containing structural and functional magnetic resonance imaging (MRI / fMRI), diffusion tensor imaging (DTI) and MEG recordings from healthy human volunteers. The MEG data is collected with a 275-channel CTF MEG system using third-order gradient balancing for noise correction and at a sampling rate of 1200 Hz with a quarter-Nyquist filter of 300 Hz. The position of the head was localized using three fiducial coils. We analyzed 6-minutes resting state (N=67) and Hariri-Hammer task (N=65) from this dataset.

The Hariri-Hammer task presents a target face, and after a short delay, two probe faces. The participant is asked to pick the probe face that matches the emotion shown in target face. As a control condition, shapes instead of faces are presented with the same design. Targets and probes are presented for 1 second. The delay between target and probe is 0.5 second. After the probe, there is a 1.5 second + jitter delay period. In total, there are 3 blocks of 30 face trials and 4 blocks of 15 shape trials. Total time to complete the task is approximately 10.3 minutes.

The preprocessing of the MEG data started with a high-pass filter at 1 Hz. Afterwards, the continuous data was segmented into 3-second epochs. Automatic artifact rejection was performed using autoreject Python package^64,65^. Autoreject looks for a threshold τ to reject epochs if the peak-to-peak amplitude (i.e. difference between lowest and highest values of signal in an epoch) is greater than τ for each channel. Briefly, it uses the following standard procedure^65^: for each candidate *τ*_*i*_, split the data in K folds. Each of the K folds along the epoch dimension (so that if there are N epochs in total, there are N/K epochs in each fold) are used as a test set once and the other K-1 folds are used as training. For each test fold, candidate threshold*τ*_*i*_ is used to reject epochs that have greater peak-to-peak amplitude than *τ*_*i*_. For each time-point, the signal is averaged over the good (i.e. not-rejected) epochs in the training set and the median is taken across all epochs in the test set. For test fold k, the Frobenius norm of the difference between mean across good epochs in training and median across all epochs in test is denoted as *ε*_*k*_. The candidate *τ*_*i*_ with the lowest average *ε* (averaged across folds) is chosen as the rejection threshold. For a bad epoch in a sensor, there can be two strategies to proceed, the epoch can be interpolated based on other channels, or if too many channels are bad, the epoch should be rejected. The method picks two parameters: κ: maximum number of bad channels for an epoch (if the number of bad channels is higher than κ, the epoch will be rejected, otherwise it will be interpolated), and ρ: the maximum number of channels that can be interpolated. For each bad channel per bad epoch, the “badness” of the channel can be quantified as the peak-to-peak amplitude. After ranking the channels from worse to best depending on this quantification, the worst ρ channels are interpolated using spherical harmonics^66^. After interpolation and rejection, the same error metric ɛ averaged across folds is used to determine the best ρ and κ using a grid search. Autoreject method is validated by comparisons with manual artifact rejection by human experts and outperformed other automatic rejection methods^65^ and is widely used in EEG/MEG literature (see for example^67–69^ among others).

After autoreject with rejection / interpolation, independent component analysis was performed with 20 components. Components that show ocular or cardiac artifacts based on a visual inspection were rejected^70,71^ and the raw data (before autoreject) was transformed with the ICAs that weren’t rejected. A 100 Hz low-pass filter and a notch filter at 60 Hz were applied. Finally, one more run of autoreject was performed on data to reject / interpolate bad epochs.

### Spatiotemporal Permutation Testing

In order to identify the channels that respond to the task, we compared event-related fields between target and control conditions using cluster-based permutation tests. The cluster-based permutation test is performed as follows: A test-statistic (F-value) is performed for every sample (across channels and time points) to compare conditions. We performed repeated measures ANOVA with encode and probe conditions as factor 1 and happy faces, sad faces and shapes as factor 2. Thresholding is performed at this stage to keep only the F-values that have a significance of p<0.001. Note that this is not the significance that results from the whole process, just a step to estimate clusters. Spatially connected F-values that were survived from thresholding are considered as clusters. The sum of these F-values in each cluster is denoted as cluster-level statistic. The cluster with the largest total F-value is selected. To calculate significance probability, Monte Carlo methods are used. Specifically, trials across two conditions are concatenated. From this concatenated set of trials, random samplings are performed N times where N is the number of trials in condition 1. F-values are calculated from these random trials and remaining trials by performing repeated measures ANOVA for each sample. Following what is done in original data, thresholding and summing F-values in survived clusters are performed. This procedure is repeated 5000 times to form the null distribution of the test statistic (i.e. maximum value of total F-value across clusters). Proportion of Monte Carlo test statistics that are larger than the observed test statistic is the significance probability, also known as p-value.

### Bayesian Regression Models

We used hierarchical Bayesian models to estimate regression coefficients using PyMC^72,73^, ArViz^74^ and Bambi^75^. We fit the models using Hamiltonian Monte-Carlo sampling with NUTS method^76^. Convergence diagnostics were assessed using posterior predictive checks, trace plots, Rubin – Gelman statistic *r*^^77^ and effective sample sizes. Monte-Carlo sampling was performed with 4 chains with 4000 tuning and 1000 inference draws for each chain. Acceptance rate is set to 0.99. Inference diagnostics (trace plots, ArViz summaries, prior and posterior checks) as well as draws can be found in the open repository osf.io/exck3^78^.

All *r*^ values were below 1.01. All effective sample sizes were larger than 300. In addition to posterior distributions for regression coefficients and HDI values, we also provide probability of direction (pd) and percentage of the posterior lying outside the region of practical equivalence (ROPE). The probability of direction is the percentage of posterior distribution that has the same sign as its mean. Therefore it is numerically very similar to the frequentist p-values and enables comparison with them. Specifically, for a two-sided p-value, *p* = 2(1 − *pd*)^79^. For the ROPE calculations, we set the limits of region of practical equivalence from −0.1 to 0.1 after standardizing the variables via taking z-scores^80^.

The models indicated in figure 5 are of the following forms:

Model 1:

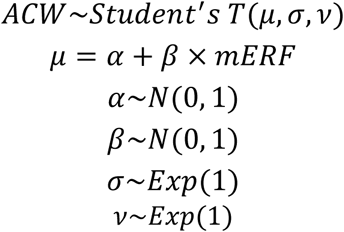

Model 2:

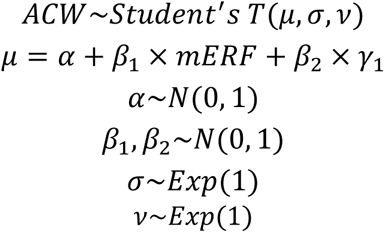

Adding *γ*_1_ to the model stratifies the relationship by *γ*_1_, letting us see if the original correlation is influenced by *γ*_1_.

The model in figure 9 investigates the relationship between ACW and mERF. We used a hierarchical model with fixed effects for mERF and random effects for clusters, trials and channels and dependent variable as ACW. We used non-centered reparametrization for channels, clusters and trials.

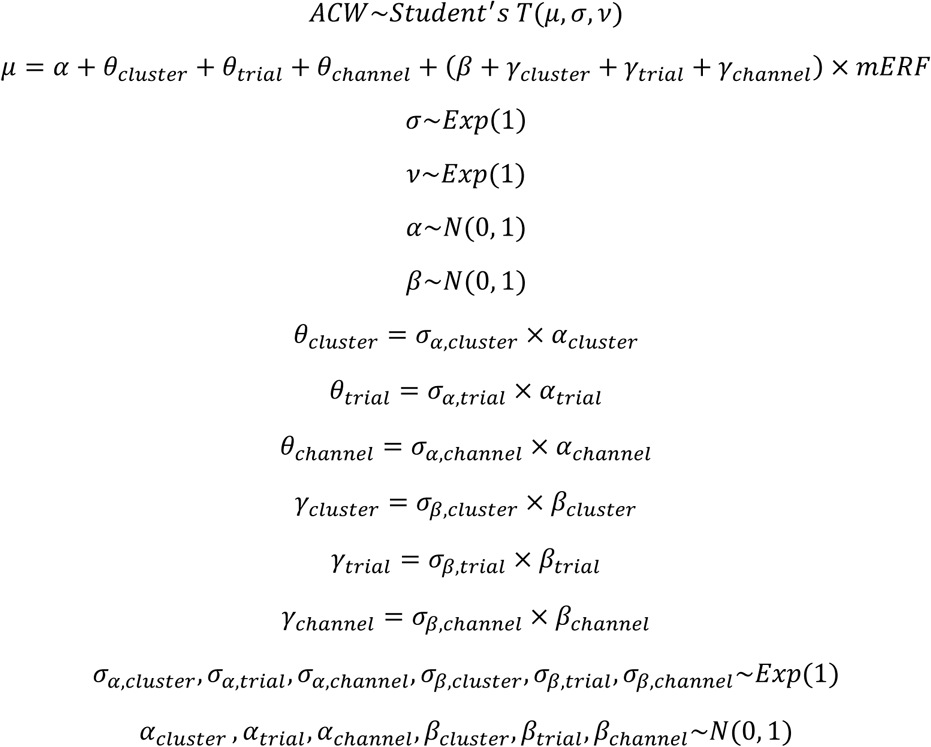

In the supplementary figures, we also show hierarchical regression models with reaction times as dependent variable and ACW as independent variable (in supplementary figure 6) and mERF as independent variable (supplementary figures 5).

### Data and Code Availability

The MEG data used in this paper is available at https://doi.org/10.18112/openneuro.ds004215.v1.0.0. The code to replicate analyses is available at github.com/duodenum96/erf_acw2.

## Results

### Computational Modeling I – Identifying a Common Neurobiological Basis for Event-Related Field Magnitudes and Intrinsic Neural Timescales

We started our inquiry by identifying a common neurobiological basis that influences both magnitudes of event-related fields (mERFs) and intrinsic neural timescales (INTs) using the Jansen-Rit model, a neural mass model that generates event-related fields (ERFs) when stimulated with an external input.

The Jansen-Rit model contains three subpopulations for every cortical column: excitatory interneurons, inhibitory interneurons and pyramidal neurons. These populations are coupled to each other via excitatory (*γ*_1_, *γ*_2_, *γ*_3_) and inhibitory connections (*γ*_4_) (figure 2a). We used a model where two of these populations are coupled to each other hierarchically via simplified Felleman and Van Essen rules^50,81^. Pyramidal neurons in a region that is lower in the hierarchy (region 1) sends feedforward connections (*A*_*F*_) to excitatory interneurons in the hierarchically upper region (region 2). Pyramidal neurons of region 2 in return, sends backwards connections (*A*_*B*_) to pyramidal cells and inhibitory interneurons of region 1. The pyramidal neurons of both regions are connected to every subpopulation of the other ROI via lateral connections *A*_*L*_^50^ (figure 2b). In total, the parameters *γ*_*i*_ and *A*_*i*_ define our set of structural parameters. Values for *γ*_2,3,4_ are empirically constrained by *γ*_1_ with relations *γ*_2_ = 0.8*γ*_1_ and *γ*_3_ = *γ*_4_ = 0.25*γ*_1_, hence, we only changed the values for *γ*_1_ and determined *γ*_2,3,4_ according to the aforementioned relations.

**Figure 2.**
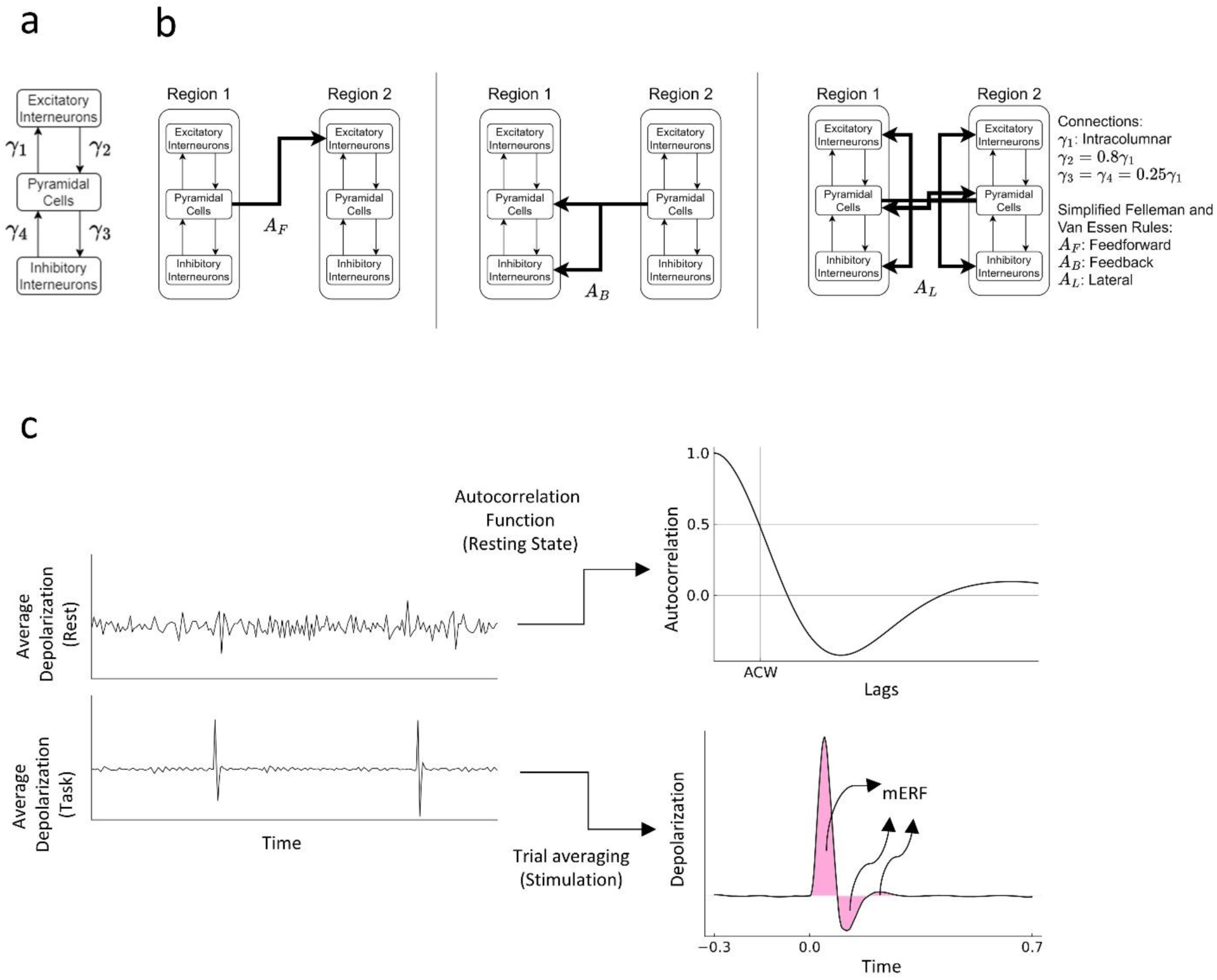
Jansen-Rit model. a. Schema of one cortical column in the Jansen-Rit model. b. Schema of coupled cortical columns and external connections between them. c. An example time series of both rest and task states as well as exemplar autocorrelation function and event-related fields. We calculated the autocorrelation window (ACW) as the lag where the autocorrelation function reaches 0.5. magnitude of event-related fields (mERFs) were calculated as the root mean squared value of the ERF after the stimulus onset.

To estimate the output measures, we did two batches of simulations corresponding to rest and task states. For the resting state, we simulated the Jansen-Rit model for 100 seconds and calculated the ACW defined as the lag where autocorrelation function reaches 0.5 in every 10 second non-overlapping window. We averaged these ACWs to get a less noisy ACW in every simulation akin to Honey et al.^82^. For the task state, we stimulated the hierarchically lower region with a delta function every 5 seconds. The period from 0.3 seconds before the stimulation to 0.7 seconds after it defines a trial. To estimate the ERFs, we averaged the depolarization of each region across trials which gives us ERFs of the form widely observed in empirical data. We parametrized the ERFs by calculating the root-mean squared value of the ERFs after the trial onset, giving us magnitude of ERFs (mERFs).

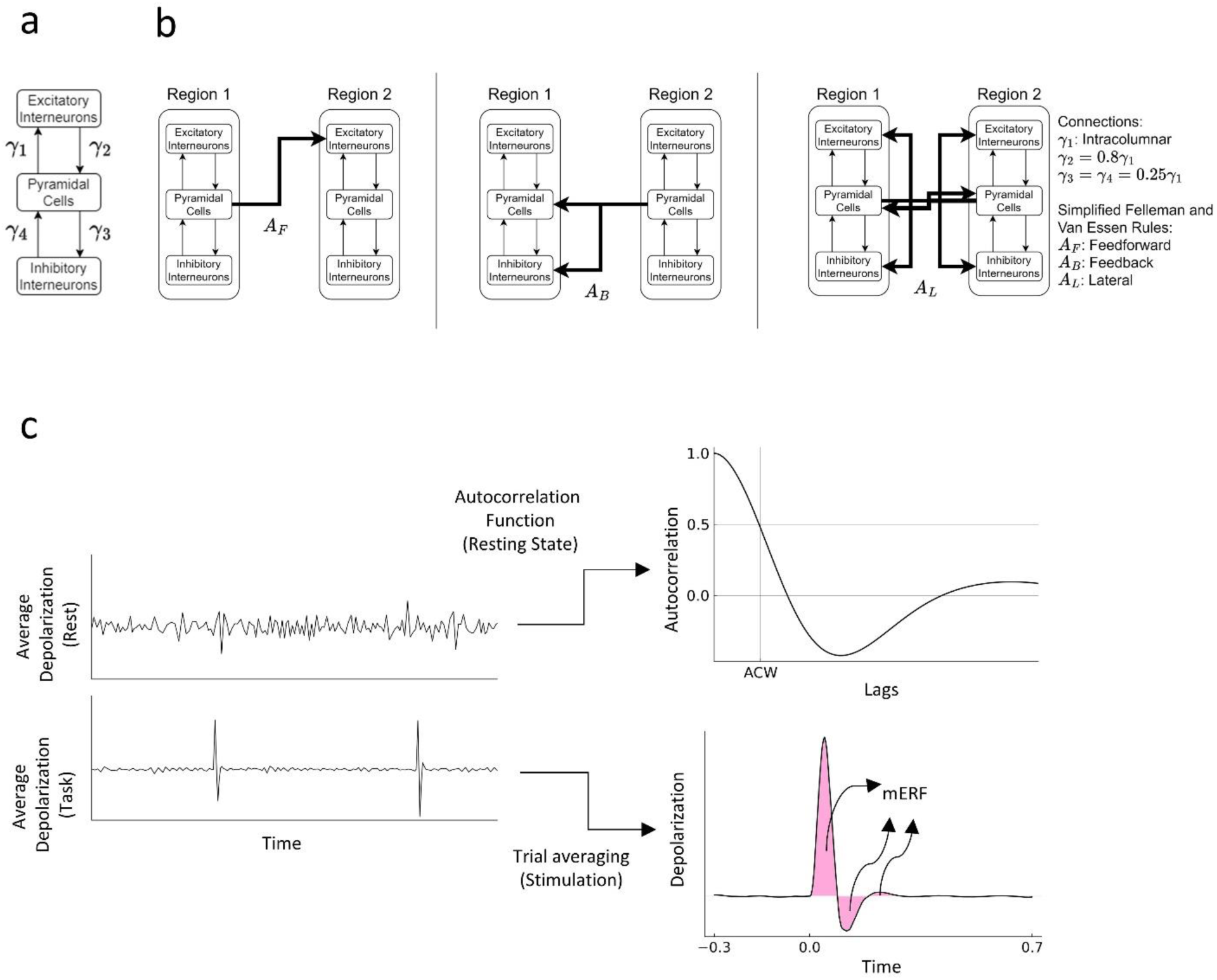

Figure c shows an example time series from rest and task conditions, autocorrelation function and event-related fields.

In Figure 3, we present the Spearman correlation between structural parameters (intracolumnar connections *γ* and long-range connections between columns (*A*_*i*_) and mERF/ACW estimates. For each parameter, we fixed all the other parameters constant and sweeped a range for the remaining parameter (40 to 70, in 31 steps for *γ*_1_, 0 to 20 in 31 steps for *A*_*i*_). For each value of the parameter, we performed 20 simulations. Our analyses shows that only the intracolumnar connections (*γ*) correlate significantly with both mERF and INT (area 1 ACW: r(615) = 0.976, p<0.001; area 2 ACW: r(615) = 0.950, p<0.001; area 1 mERF: r(615) = 0.928, p<0.001; area 2 mERF: r(615) = 0.976, p<0.001). In contrast, feedforward connections (*A*_*F*_) only correlates with the mERF of the second area since it conveys the input from first area to second area (area 1 ACW: r(615) = 0.034, p=1; area 2 ACW: r(615) =-0.02, p=0.623; area 1 mERF: r(615) = −0.045, p=0.269; area 2 mERF: r(615) = 0.988, p<0.001). Lateral connections (*A*_*L*_) on the other hand are inversely correlated with the INT of both areas (area 1 ACW: r(615) = −0.842, p<0.001; area 2 ACW: r(615) =-0.858, p<0.001; area 1 mERF: r(615) = 0.054, p=1; area 2 mERF: r(615) = 0.064, p=1). Finally, backward connections (*A*_*B*_) are positively correlated with mERF in both areas and negatively correlated with the ACW of only the first area (area 1 ACW: r(615) =-0.535, p<0.001; area 2 ACW: r(615) = −0.096, p=0.274; area 1 mERF: r(615) =0.410, p<0.001; area 2 mERF: r(615) = 0.993, p<0.001).

**Figure 3.**
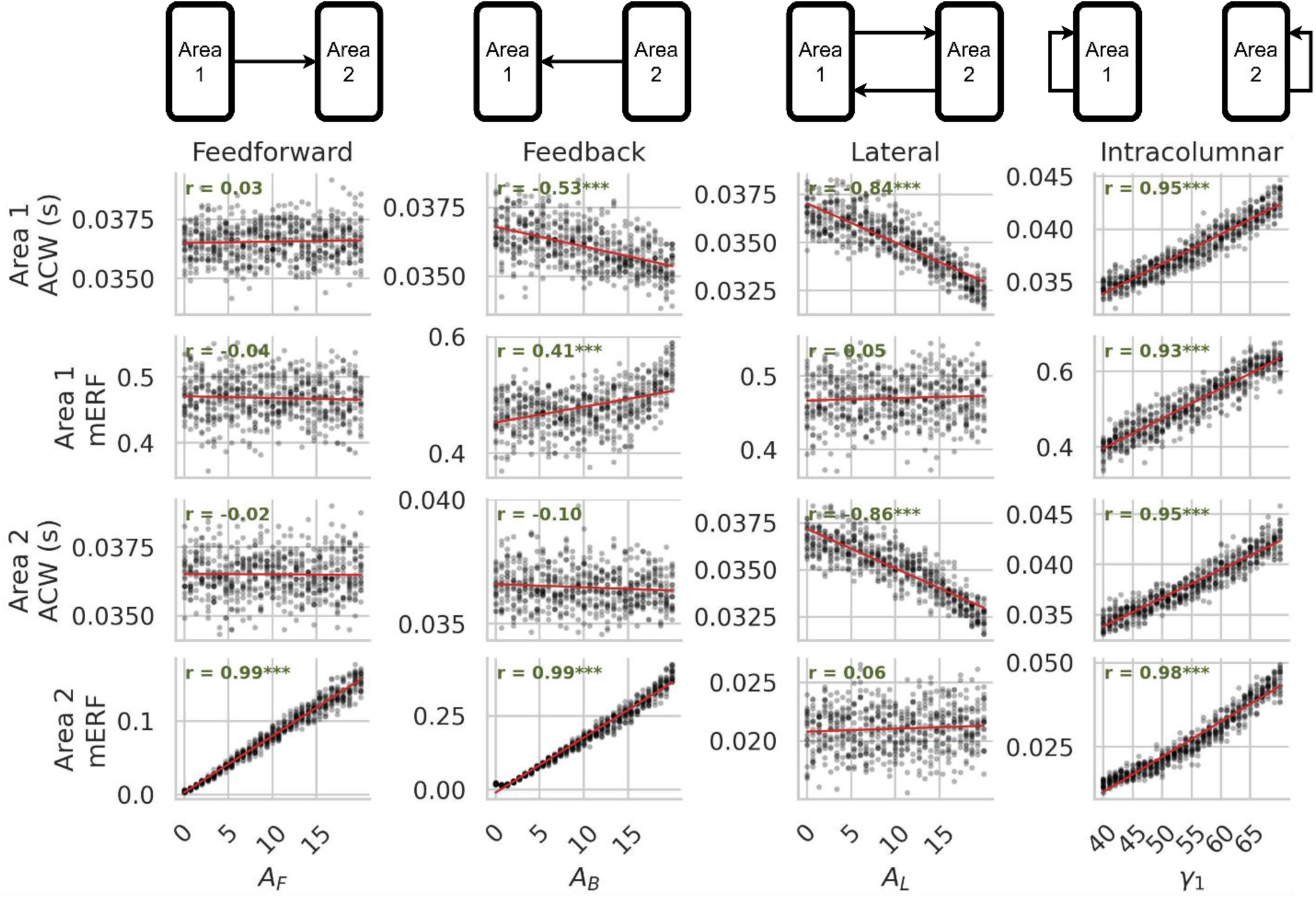
Identifying a potential common neurobiological basis for magnitude of event-related fields (mERFs) and autocorrelation windows (ACWs). For each model parameter, every other parameter was fixed and we swept across different values of the parameter, performing 20 simulations in rest (for ACW) and task (for mERF) conditions. We evaluated the relationship between model parameters and mERF/ACW via Spearman correlations. Asterisks indicate statistical significance after Bonferroni-Holmes correction (*: p<0.05, **: p<0.01, ***: p<0.001).

Our results indicate that both outcome variables, ACW and mERF, are both correlated only to intracolumnar/intra-regional excitatory and inhibitory connections suggesting that the latter provides a shared biological-structural substrate for both ACW and mERF. In contrast, inter-regional connections are not shared by ACW and mERF but influence the two measures in differential ways. Specifically, lateral connections seem to influence only the ACW but not the mERF while both feedbforward and feedback connections more strongly impact the mERF (and less so the ACW). Summing up, intra-columnar connections are shared by both ACW and mERF while inter-regional connections (lateral, feedforward and feedback) exert differential impact on the two measures. Given that our aim was to identify a common substrate for ACW and mERF, we will focus on the intracolumnar connections. Hence, in the next section, we explore this dependence in greater detail by simplifying the model to a single region and systematically varying its intracolumnar connections to assess the relationship between mERF and ACW.

### Computational Modeling II – Relationship Between Intracolumnar Connections, Intrinsic Neural Timescales, and Event-related Fields in the Jansen-Rit Model

Building on the finding that both ACW and mERF are related to changes in the model’s intracolumnar connections, we next examined how these outcome measures depend on each other for different values of these intracolumnar connections. Specifically, we analyzed the relationships between ACW, mERF, and intracolumnar connections (*γ*_1_) by simulating the model with different values of *γ*_1_. Since our focus is solely on intracolumnar connections, we conducted these simulations using a single region. As in previous simulations, we assessed resting state ACW by calculating it in 10-second non-overlapping windows, and estimated mERF from trials during the task state. We varied *γ*_1_ within a range of 40 to 70, sweeping in increments of 1 to cover a broad parameter space. For each *γ*_1_ value, we performed 20 simulations. As in the section above, the values for other intracolumnar connections *γ*_2,3,4_ were determined by empirical constraints *γ*_2_=0.8*γ*_1_ and *γ*_3_=*γ*_4_=0.25*γ*_1_, thus changing the value of *γ*_1_ scales both excitatory (*γ*_1,2,3_) and inhibitory (*γ*_4_) intracolumnar connections.

In Figure 4b, we report the correlations between both ACW and mERF with intracolumnar connections (*γ*_1_). We find a positive correlation between *γ*_1_ and ACW (r(420) = 0.784, p < 0.001), as well as a stronger positive correlation between *γ*_1_ and mERF (r(420) = 0.904, p < 0.001). Figure 4c presents the correlation between ACW and mERF, which shows a moderate positive relationship (r(420) = 0.517, p < 0.001). The positive correlation can be framed with the fluctuation-dissipation theorem^44,46^ positing that the temporal structure of a system’s spontaneous activity predicts its response to external inputs. In the supplementary material, we give a brief and self-consistent introduction to this theorem in the context of event-related brain activity. These findings further support the close relationship of intra-columnar connections with both ACW and mERF including the latter two’s association.

**Figure 4.**
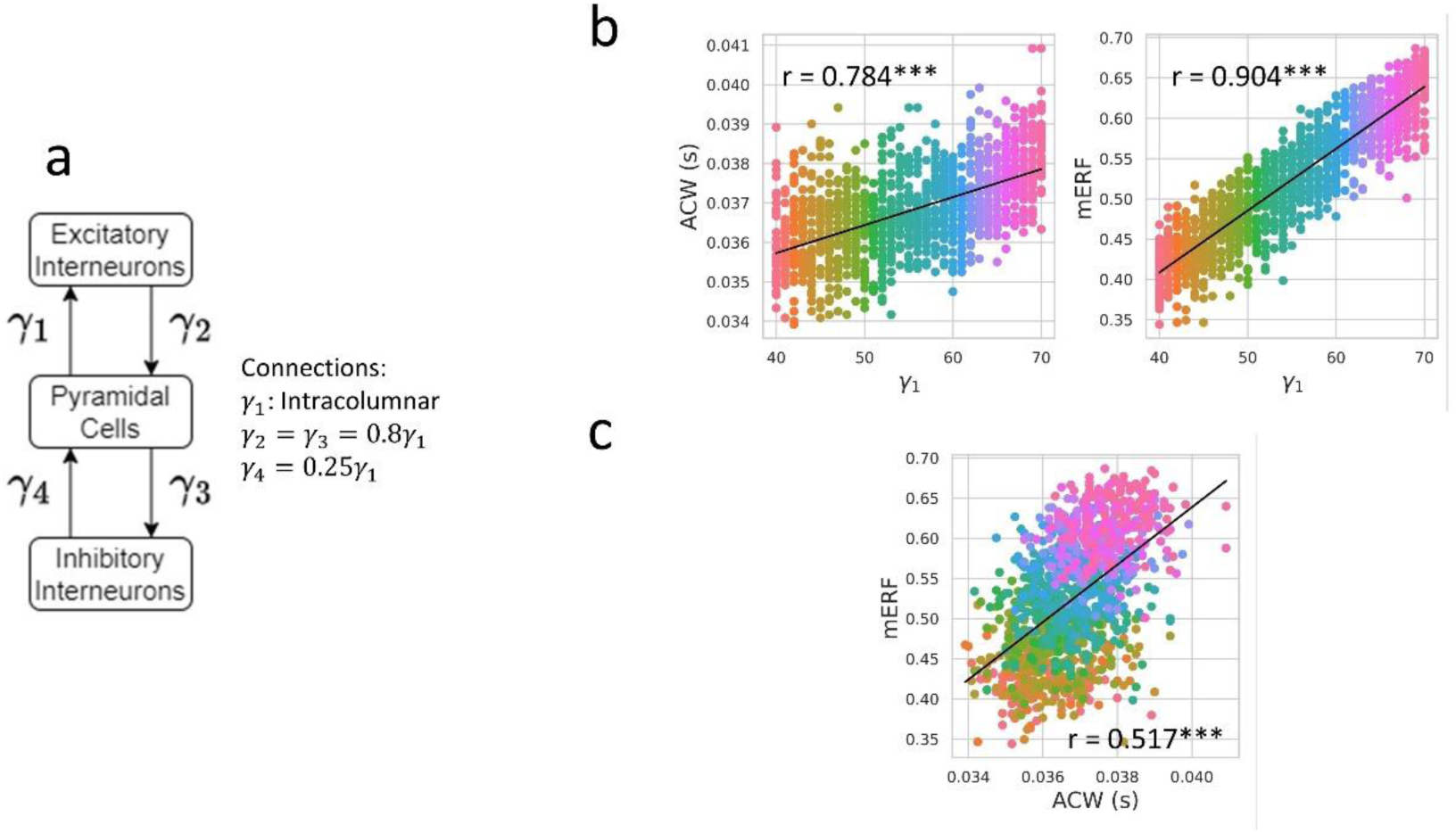
Relationships between intracolumnar connections (*γ*_1_), autocorrelation window (ACW), magnitude of event-related fields (mERF). b. Correlation between dynamic variables and intracolumnar connections (*γ*_1_) c. Correlations between dynamic variables (ACW, mERF) with each other. Colors denote different values of *γ*_1_.

Following these results, we next hypothesized that the correlation between ACW and mERF depends on the strength of intracolumnar connections as it is the only parameter systematically varied across our simulations. To test this hypothesis, we fit two regression models. In the first model, mERF is predicted solely by ACW. In the second model, mERF is predicted by both ACW and intracolumnar connections (*γ*_1_). If our hypothesis is correct, we expect to observe a reduction in the regression coefficient between ACW and mERF when intracolumnar connections are included to multivariate regression, as the variance in mERF that is explained by ACW should be largely accounted for by the intracolumnar connections (*γ*_1_).

To test these relationships, we employed Bayesian regression models. The unconditioned model is represented as *mERF* = *βACW*, while the conditioned model is expressed as *mERF* = *β*_1_ *ACW* + *β*_2_*γ*_1_. To estimate the posterior distributions of the regression coefficients for each model, we utilized Monte Carlo sampling via the No-U-Turn Sampler (NUTS) method^76^, implemented in pyMC^72,73^. Detailed model specifications and priors are provided in the Methods section. Traces, trace plots, ArViz summaries showing mean, standard deviation, 94% HDI, effective sample size, Rubin-Gelman statistic *r*^, probability of direction (pd) and percentage in the region of practical equivalence (ROPE) for each parameter, prior and posterior predictive checks for each parameter are available in the open data repository^87^. For ease of interpretation, all variables were standardized using z-scores. The proportion of the posterior distribution inside 94% HDI that shares the same sign as its mean is referred to as pd, which is related to the frequentist two-sided p-value by the formula p=2(1-pd). The pd value ranges between 0 and 1, with higher values indicating greater certainty about the existence of an effect. The percentage in ROPE represents the proportion of the posterior distribution inside 94% HDI that lies within a predefined range, where results can be considered equivalent to no effect. Following Kruschke^80^, we define this range as [−0.1, 0.1]. A lower percentage within the ROPE indicates stronger evidence against the null hypothesis, suggesting that the result cannot be considered equivalent to zero.

Figure 5a displays the estimated regression coefficients along with their 94% HDIs. The results of the unconditioned model indicate a significant positive prediction of ACW by mERF (mean: 0.524, SD: 0.024, 94% HDI: [0.478, 0.567], pd: 1, % in ROPE: 0). As anticipated, conditioning the relationship between ACW and mERF on *γ*_1_ resulted in the shrinkage of the regression coefficient for mERF (mean: −0.026, SD: 0.054, 94% HDI: [-0.12, 0.083], pd: 0.686, % in ROPE: 0.957) and a substantial positive coefficient for *γ*_1_ (mean: 0.609, SD: 0.054, HDI: [0.504, 0.705], pd: 1, % in ROPE: 0). In addition to the multiple regression analysis, we also present correlation coefficients between mERF and ACW for fixed values of *γ*_1_ in Figure 5b. These results show that the correlation between mERF and ACW diminishes as when *γ*_1_ is fixed (*γ*_1_=40: r(420) = −0.058, p=0.719; *γ*_1_=50: r(420) = 0.096, p=0.719; *γ*_1_=60: r(420) = −0.278, p=0.248).

**Figure 5.**
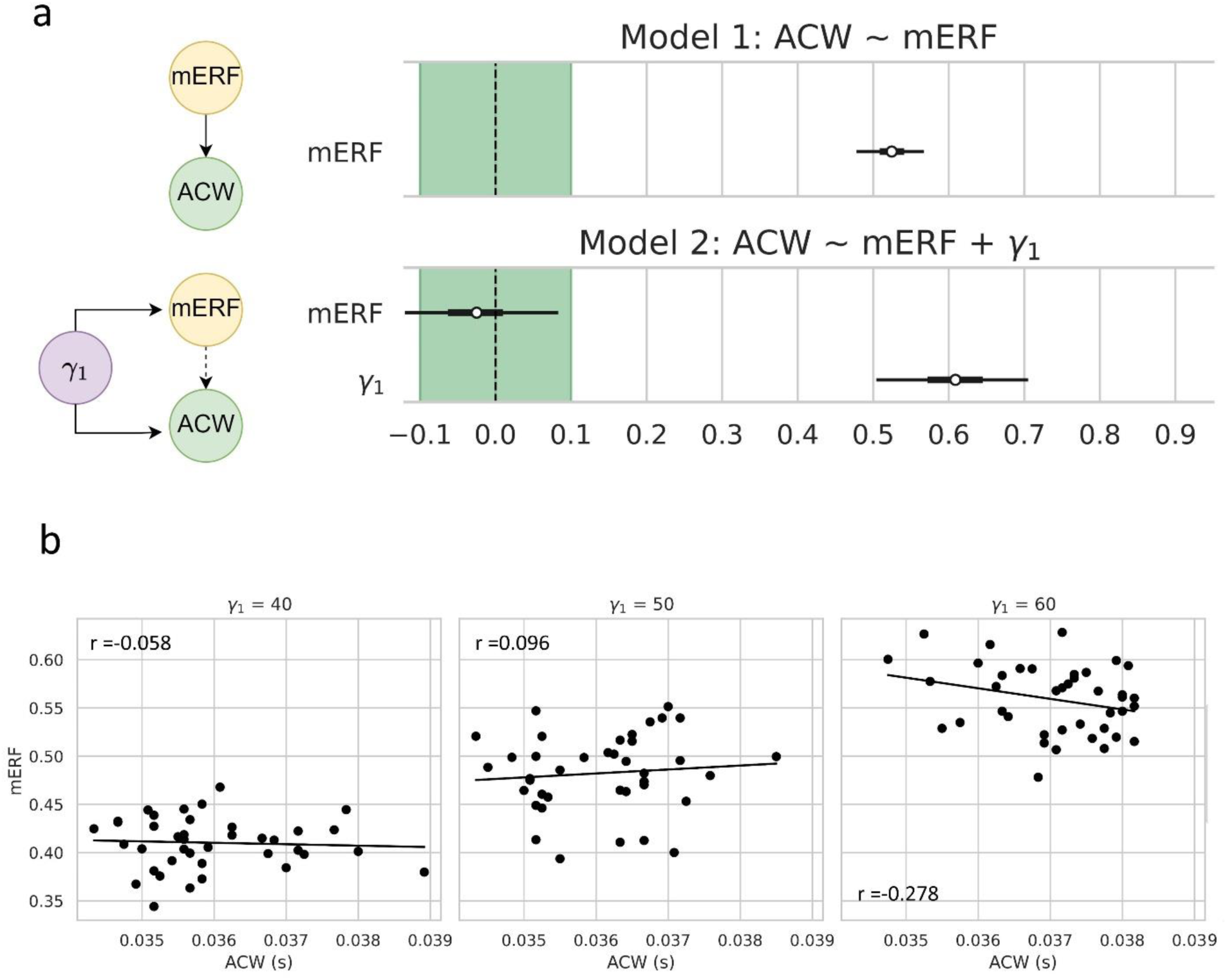
The relationship between ACW and mERF depends on *γ*_1_. a. On top, we see the forest plot that shows the posterior regression coefficient of the slope in a Bayesian linear regression between mERF and ACW. The bottom panel shows the multivariate regression results where *γ*_1_ is introduced as an additional variable. The shrinkage of the coefficient for ACW indicates the relationship between ACW and mERF depend on *γ*_1_. b. Scatter plots where we fix *γ*_1_ and inspect that the ACW-mERF correlation disappears when *γ*_1_ is fixed.

In summary, these results provide valuable insights into the relationships between intrinsic neural timescales (INTs), magnitude of event-related fields (mERFs), and their dependence on intracolumnar connections. A positive correlation between mERFs and autocorrelation windows (ACWs) was observed, and this relationship was shown to be modulated by the underlying structural variable, intracolumnar excitatory and inhibitory connections (as distinguished from feedforward, feedback and lateral connections between the columns of different regions). In the following section, we turn to an empirical dataset to assess how these computational findings align with observed data.

### Empirical Data I - Event-related Correlates of Emotional Face Recognition

In the second leg of our investigation, we analyzed data from the NIMH Intramural Healthy Volunteer Study^53^, which includes magnetoencephalogram (MEG) recordings collected during both resting-state and a facial emotion recognition task (Hariri-Hammer task^83^). In the Hariri-Hammer task, participants were presented with "encode" or "probe" trials within both face and shape blocks. In the encode trials, participants viewed either a happy or sad face in the face block or a shape in the shape block. In the probe trials, two faces (in the face block) or two shapes (in the shape block) were shown, and participants were asked to identify the item that matched the one presented in the corresponding encode trial. A schematic of the experimental design is shown in Figure 6a.

**Figure 6.**
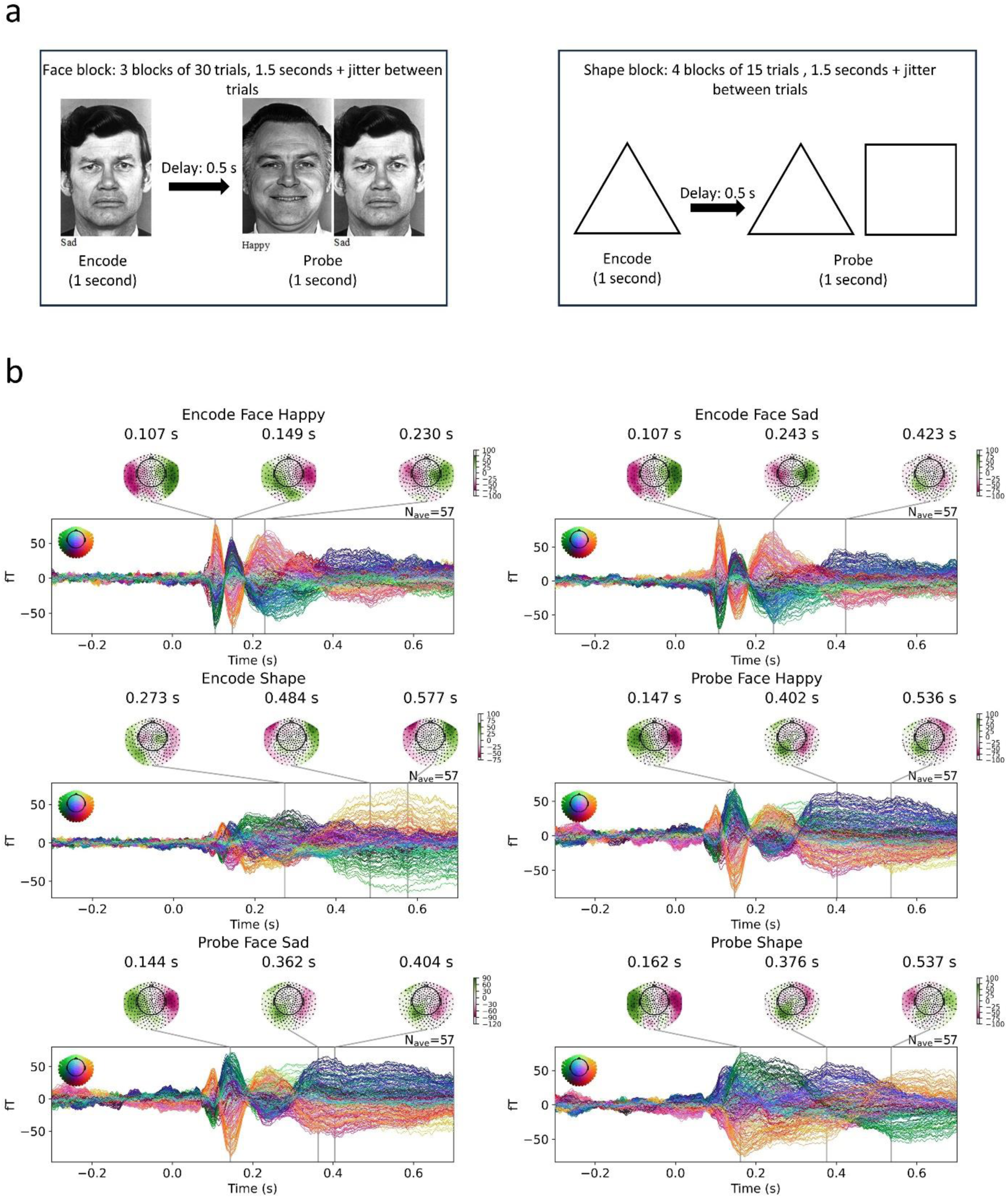
Task design of Hariri – Hammer and event-related fields. a. A schema of the task design. Hariri – Hammer task is a task designed to assess emotional face recognition and consists of face and shape blocks. In encode condition, a participant is presented with either a face or shape. In the probe condition, they are presented with two stimuli and asked to pick the one that was presented in the encode condition. b. Event-related fields for each condition and their topographic distribution at extrema of event-related fields. Each line corresponds to one channel.

We began our analysis by calculating the event-related fields (ERFs) for each condition and performing a spatiotemporal permutation test to identify clusters and time windows that exhibited the strongest contrast between conditions. Figure 6b presents the ERFs corresponding to each experimental condition. A repeated measures analysis of variance (rmANOVA) was conducted where one factor is the moment (encode versus probe) and the other is the content (happy faces, sad faces and shapes).

Figure 7 presents the results of the spatiotemporal permutation test for the factors of happy face, sad face, and shape. Six spatiotemporal clusters were identified that exhibited significant differences between the three conditions. These clusters were primarily located in the left and right temporal sensor regions. In the encode condition, two primary temporal clusters were found: one spanning the 0.074 to 0.122–0.124 second range and another from 0.163–0.170 to 0.347–0.372 seconds following stimulus onset. In the probe condition, a significant difference was observed only within the 0.074 to 0.122–0.124 second range.

**Figure 7.**
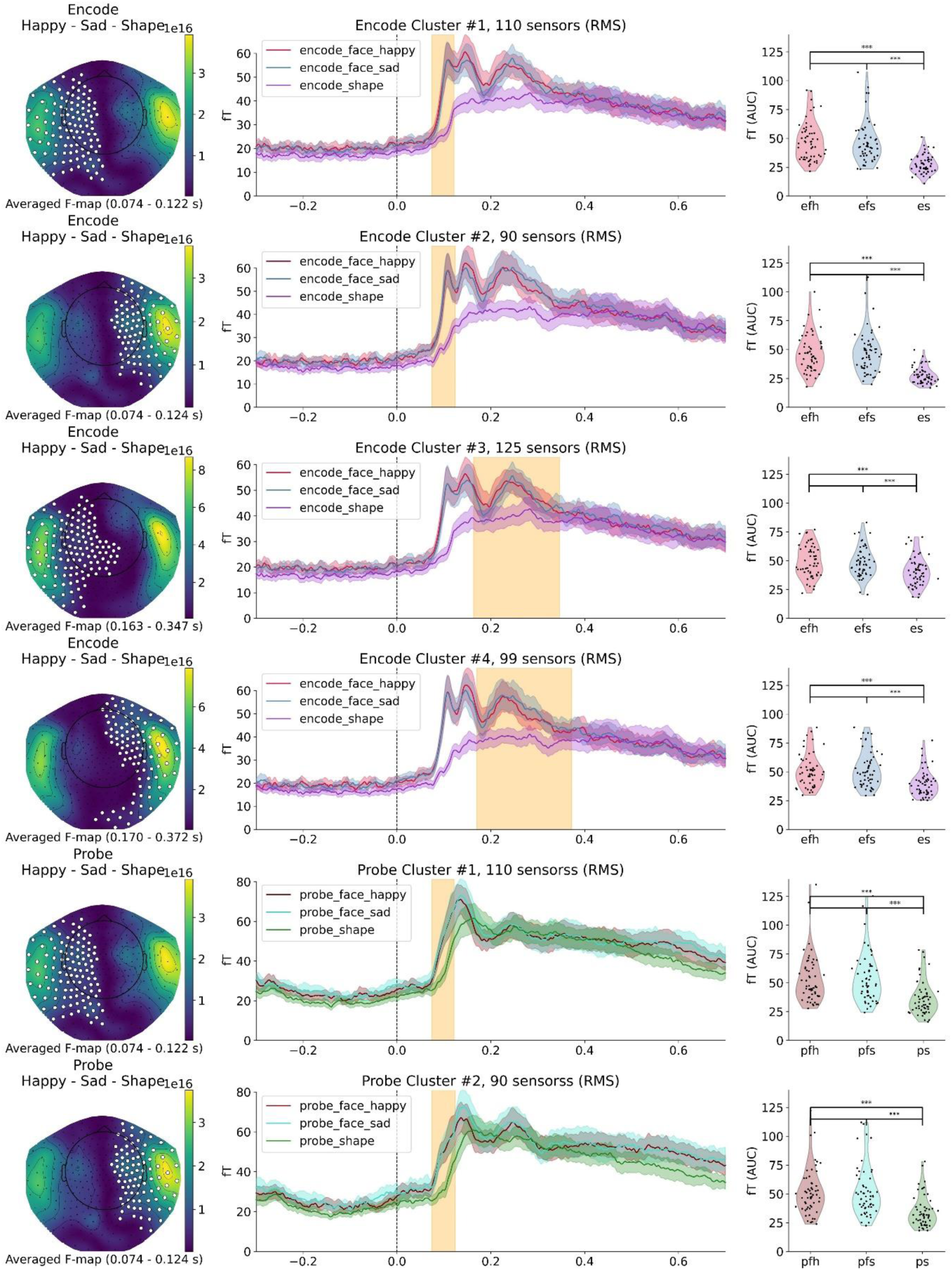
The results of the spatiotemporal permutation test. The first column shows the spatial cluster that showed a significant difference between conditions. Black dots denote channels and white dots denote cluster. Color corresponds to the F-statistic from the repeated measures ANOVA test. The second column shows the time course of the root-mean squared value of the event-related field (ERF). The temporal cluster that was found to show significant difference between conditions corresponding to the spatial cluster on the left is shown with yellow vertical coloring. The shades around event-related fields show the standard deviation of event-related fields. In the third column, we compare the average value of the root-mean squared ERF values between three conditions. Asterisks denote significance.

We then performed multiple comparisons on the root-mean squared (mERFs) averaged over time for the three conditions. The results of these comparisons, along with the corrected p-values for each test, are provided in Supplementary Table 1. Notably, while a significant difference was observed between the face and shape conditions, no statistically significant difference was found between the happy face and sad face conditions. Only clusters that exhibited significant differences following multiple comparisons are presented in the main paper. Clusters that did not show significant differences after correction can be found in Supplementary Figures 1 to 4. In summary, we identified significant differences in mERFs between the face and shape conditions in temporal sensors, primarily occurring at 100 ms and 250 ms post-stimulus onset.

### Empirical Data II - Relationship Between Intrinsic Neural Timescales and Event-Related Fields

The next step of our analysis focuses on exploring the relationship between intrinsic neural timescales (INTs) and event-related fields (ERFs). To estimate INTs, we calculated the autocorrelation window (ACW), defined as the lag at which the autocorrelation function reaches a value of 0.5^82^, using the preprocessed resting state data from our participants. As in the simulations, we computed one ACW value for each 10-second, non-overlapping time window and averaged those values to obtain a single ACW estimate per channel for each subject. Figure 8a displays the autocorrelation functions for each subject, averaged across channels. Figure 8b illustrates the distribution of ACW values across the scalp, averaged across subjects.

**Figure 8.**
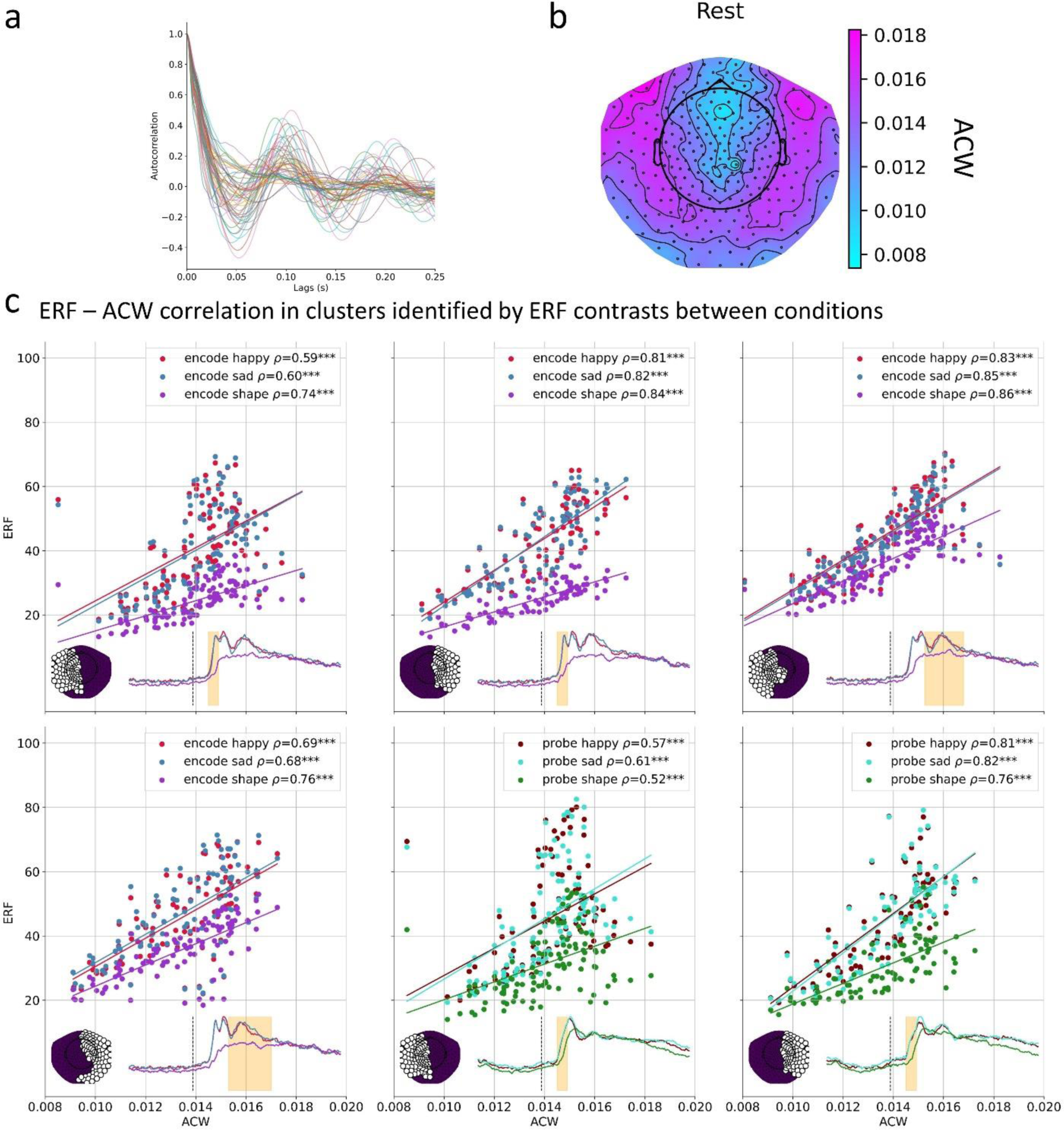
a. Autocorrelation function of all subjects averaged across channels. b. Topoplot of autocorrelation window (ACW) across the scalp. We calculated ACW as the lag where autocorrelation function reaches 0.5 for each 10 second non-overlapping window from resting state scans. We averaged those ACW values to obtain one ACW estimate per subject per channel. c. Correlation between mERF and ACW for each identified spatiotemporal cluster.

In Figure 8b, we show the positive relationship between ACW and mERF values for each spatiotemporal cluster, consistent with the results observed in the modeling. Given the nested structure of our data— where mERFs are nested within clusters (identified through spatiotemporal permutation testing), trial types (e.g., encode happy face, probe shape), and MEG channels—we employed Bayesian hierarchical models for analysis. We employed a hierarchical Bayesian model with varying intercepts and varying slopes to account for the nested structure of our data. In this model, the dependent variable is ACW, and we included fixed effects for both the intercept and slope of mERF. Additionally, we modeled random effects for intercepts and slopes at the level of clusters (obtained via spatiotemporal permutation testing), trial types (e.g. encode face happy, probe shape etc.), and MEG channels. This hierarchical structure allows for the modeling of both specific and group-level variations, thereby facilitating a more nuanced understanding of the relationships between ACW and mERF across different levels of the data hierarchy. By incorporating random effects for clusters, trial types and channels, the model accounts for the variability within each level, while still pooling information across all data points, which enhances the precision of parameter estimates and improves the generalizability of the findings^84^.

Figure 9a shows the regression coefficient estimates after z-score standardization. We observe a positive fixed effect for the slope of mERF, which aligns with the findings from our modeling results (mean: 0.224, SD: 0.034, 94% HDI: [0.163, 0.289], pd: 1, ROPE: 0). This effect remains robust across variations in clusters and trial types, as indicated by the ROPE values (>0.95 for all clusters and trial types; full results and diagnostic checks are provided in the open repository^87^ as usual). To assess whether the positive relationship between mERF and ACW holds across all clusters and trial types, we combined the posterior samples from the fixed effect and the random effects. Specifically, we added the posterior samples of the fixed effect and random effects to construct new coefficients that reflect both sources of variation. The resulting coefficients were then summarized by calculating the mean, standard deviation (SD), and the highest density interval (HDI) of the combined posterior distribution. This approach allows us to account for both the fixed and random effects when evaluating the overall relationship between mERF and ACW. The analysis confirms that the positive relationship between mERF and ACW is consistent and robust across clusters and trial types (Figure 9a, denoted as ERF+ERF|Cluster and ERF+ERF|Trial for clusters and trial types, respectively; pd=1 and ROPE=0 for all clusters and trial types).

**Figure 9.**
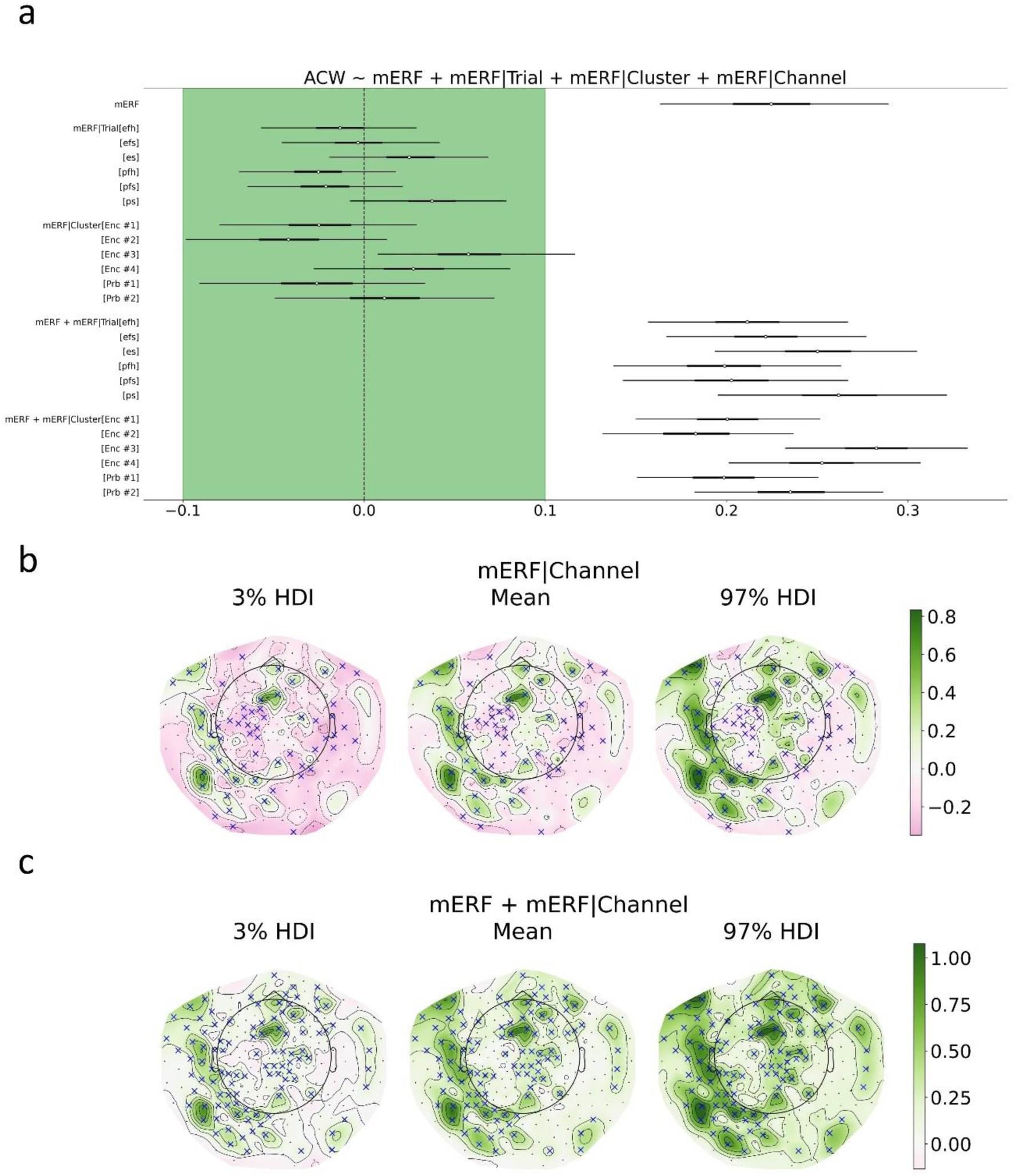
Relationship between intrinsic neural timescales and event-related fields. a. Coefficient estimates for a hierarchical Bayesian model where ACW depends on mERF nested inside cluster, trial and channel variables. Dots denote the averages and lines denote 94% highest density probability intervals (HDI) of posterior distributions for coefficients. Limits for Region of Practical Equivalence (ROPE) is indicated with the green shading. b. Channel specific slopes estimated in the hierarchical model. c. Total effect for channels as a sum of fixed effect and random effect for each channel. In panels b and c, channels that have a ROPE less than 0.01 are indicated with a blue cross. Abbreviations: efh: encode face happy, efs: encode face sad, es: encode shape, pfh: probe face happy, pfs: probe face sad, ps: probe shape, Enc: Encode, Prb: Probe.

In Figure 9b, we plot the mean and 94% HDI values for the channel-specific random effects. Channels with a ROPE greater than 0.99 are marked with blue crosses. The total effect for each channel, obtained by adding the samples from both the fixed and random effects as described previously, is shown in Figure 9c. These results further support the positive relationship of mERF and ACW, as evidenced by the overall positive effect, with the only channels exhibiting a ROPE greater than 0.99 being those with a positive effect.

In summary, our empirical results replicate and thus validate the positive relationship between ACW and mERF as predicted by the Jansen-Rit model of cortical columns. Additionally, we investigated the relationships between ACW and reaction times, as well as mERF and reaction times. The detailed results of these analyses can be found in the supplementary figures 5 and 6.

## Discussion

In this study, we aimed to investigate the relationship between intrinsic neural timescales (INTs) during the resting state and event-related activity during the task state by searching for their shared underlying structural-computational substrate. We employed computational modeling, which was complemented by the analysis of magnetoencephalogram (MEG) data. The Jansen-Rit model used for event-related field (ERF) generation demonstrated that the relationship between INTs and ERFs is strongly influenced by intracolumnar/intraregional connections rather than inter-regional connections between different columns: higher intracolumnar excitatory and inhibitory connectivity resulted in both increased ERF magnitudes (mERF) and longer autocorrelation window (ACW) values. Both the computational model and the empirical data revealed a positive relationship between ACW and ERF. These findings suggest that intracolumnar connections play a critical role in mediating both resting-state ACWs and task-related ERFs, highlighting a close connection between the brain’s intrinsic dynamics and its task-specific responses to external stimuli.

### Computational insights – intracolumnar connections influence both INT and Event-related Activity

Our computational analysis demonstrated that both the autocorrelation window (ACW) and the magnitude of event-related fields (mERF) are both sensitive only to variations in the intracolumnar connectivity parameter within the Jansen-Rit model, a widely used framework for simulating event-related neural activity. This finding aligns with prior work by Wong and Wang (2006), who, in their influential study, modeled decision-making processes in competing neural populations with common inhibitory feedback. While their model did not explicitly examine resting-state INTs, they similarly observed that increased intra-regional recurrent connections led to longer timescales. However, their analysis did not address the strength of the neural response to external stimuli and focused on reaction times of perceptual decision making. Our work extends their framework by incorporating both resting-state INTs and task-related ERFs, highlighting the role of intracolumnar connectivity in modulating both spontaneous and stimulus-evoked neural activity. Future research could build on this by extending the decision-making model proposed by Wong and Wang to incorporate the dynamics of cortical columns as modeled in the Jansen-Rit framework, linking event-related activity with perceptual decision-making processes.

In our paper, we focused on the shared biological basis of both mERF and ACW, the intracolumnar connections. In contrast, we did not further investigate the effect of inter-columnar connections between cortical columns since none of them were correlated with both mERF and ACW. However, it should be noted that lateral connections and feedback connections, while not significantly affecting mERFs, are still correlated with ACWs. The insignificance of backward connections for mERF can be hypothetically related to the scales of neural activity in lower (area 1) and higher (area 2) areas. Since area 1 is directly stimulated by the input, its event-related response is much larger than in area 2. Thus, the backward connection of area 2 to area 1 does not influence the magnitude of event-related field. On the other hand, feedback connections are very important for generation of various oscillatory phenomena in the Jansen-Rit model (we refer the reader to^50^ for a systematic investigation of the relationship between event-related activity and connections between cortical columns). mERF results for lateral connections are also consistent with previous investigations (see figure 8 in^50^). Further work out of the scope of this paper is needed for a more systematic investigation of external connections between cortical columns and ACWs.

Our findings are consistent with and extend a recent study using a whole-brain model, which demonstrated that changes in ACWs during external stimulation are strongly influenced by intraregional recurrent connections but only weakly by the overall inter-regional connectivity topology (Çatal et al., 2024). The pronounced dependence of intrinsic neural timescales (INTs) on intracolumnar connections can, metaphorically put, be conceptualized as a reverberatory process, where intracolumnar connections act as an "echo chamber" that sustains and amplifies the activity induced by the external stimulus within the cortical column of that region, effectively preserving it as a form of intra-regional memory. In our model, we observed an increased mERF in response to a delta-function input under increased intracolumnar connectivity: this increase suggests that stronger intracolumnar connections facilitate enhanced reverberatory processing, thereby amplifying the neural response to external stimuli within the cortical column of that region.

### Empirical data – Replication of the modelling data on the relationship of ACW and mERF

Confirming previous studies (REF), our spatiotemporal permutation testing revealed the strongest contrasts in spatial clusters within the temporal sensors, and temporal clusters within the 100 ms and 250 ms post-stimulus windows. While the spatial resolution of MEG is relatively limited compared to fMRI or intracranial methods for observing neural activity in specific regions, our findings align with previous research that has observed medial temporal activity, such as in the amygdala, during the Hariri-Hammer task^54–56,83^.

In accordance with the modeling results, we observed a positive relationship between intrinsic neural timescales (INTs) and event-related fields (mERFs). While INTs have been extensively studied in various cognitive and attentional contexts^5,10,19,85,86^, their relationship to event-related activity remains largely unexplored. As such, taken in a more general context, our study offers a potential starting point for understanding the interplay between resting-state and task-related brain dynamics. We view this analysis as an initial step and plan to pursue further investigations along this line of inquiry.

### Intrinsic Fluctuations Reflect Response to External Perturbations

In line with our research aims, our findings bridge a critical gap in our knowledge identified in the introduction—namely, the relationship between intrinsic neural timescales (INTs) and task-state event-related activity. By operationalizing INTs through the autocorrelation window (ACW) and modeling the relationship between INT and stimulus-evoked responses, we directly address how the brain’s intrinsic dynamics relate to external task-evoked activity in both modeling and empirical data. The observed positive relationship between ACW and mERF supports the hypothesis that intrinsic neural timescales, which have been linked to resting-state activity^3,4,87^, also reflect the brain’s response to external stimuli. This novel connection underscores the importance of considering resting-state dynamics for understanding task-related neural processes. This is consistent with the fluctuation-dissipation theorem which states that the response to an external perturbation has the same form of an intrinsic fluctuation in the absence of the external perturbation^44–46^. In the supplementary material, we provide a brief introduction to this central result. The main intuition is that the random forces that cause fluctuations in the resting state (due to synaptic noise^88^, ion channel fluctuations^89^, random thermal noise^90^, etc.) are qualitatively no different from an external perturbation in the form of an electrical stimulation in vitro or a cognitive task in vivo. Thus, the autocorrelation function of intrinsic fluctuations also contains information about the dissipation back to the baseline from a transitory (spontaneous or externally evoked) displacement of neural activity.

The framework of the fluctuation-dissipation theorem also offers a novel perspective on the conceptualization of intrinsic neural timescales (INTs) and their theoretical underpinnings. In their foundational work, Hasson et al.^6^ introduced the concept of “process memory,” which unifies short-term information storage and information processing within the context of neuronal dynamics. This framework posits that INTs serve as indices of a neuronal signal’s memory, with longer INTs reflecting an enhanced capacity of a brain region to retain information over extended durations. The fluctuation-dissipation theorem provides a potential theoretical basis for this "hierarchical process memory" by establishing an explicit theoretical link between intrinsic neuronal dynamics and responses to external perturbations. Specifically, a slower decay of the autocorrelation function in the resting state implies a prolonged decay of external perturbations, resulting in a delayed return to baseline activity. In this sense, the external perturbation is effectively retained within the "process memory" of the brain region for a longer period.

## Limitations

Several limitations of this study must be acknowledged. First, while our analysis establishes a causal relationship between the magnitude of event-related fields (mERFs) and intrinsic neural timescales (INTs) with intracolumnar connections in the computational model, we do not claim biological causality. To demonstrate causal relationships in a biological sense, future studies would need to incorporate direct measurements of intracolumnar connectivity, as well as methods such as do-calculus to appropriately randomize these connections in empirical data^91^. Second, our analysis of empirical data did not include source reconstruction, as our primary goal was not to localize specific cognitive tasks, which have been addressed in prior studies (e.g., Hariri-Hammer task^83^), but rather to investigate the dynamic relationships among ERFs, INTs, and intracolumnar connectivity. A third limitation pertains to the methodological constraints of the empirical MEG data: we were unable to directly measure or model intracolumnar connections within the data itself. Future work will need to address this by incorporating more advanced imaging techniques or multi-modal data that can probe intracolumnar dynamics more directly.

## Conclusion

In recent years, intrinsic neural timescales (INTs) have garnered significant attention, with numerous studies investigating their role in both empirical and theoretical contexts. However, the relationship between the brain’s resting-state INTs and its event-related activity remains poorly understood. To explore this relationship, we used computational modeling in combination with an analysis of magnetoencephalography (MEG) data collected during an emotional face recognition paradigm. Our findings reveal a positive relationship between the magnitude of event-related fields (mERFs) and INTs in both the computational model and the empirical data. Moreover, the modeling results suggest that the relationship between INTs and mERFs is strongly influenced by excitatory and inhibitory intracolumnar/intraregional connections. We conclude that intracolumnar connections serve as a common computational-structural substrate for both the brain’s INTs and event-related activity, highlighting their role in connecting the relationship between these two phenomena. More generally, this means that intra-columnar connections provide an intrinsic connection of the resting state’s INTs with task-related activity.

## Acknowledgements

G.N. is supported by the European Union’s Horizon 2020 Framework Program for Research and Innovation under the Specific Grant Agreement no. 785907 (Human Brain Project SGA2), UMRF, uOBMRI, CIHR, and PSI. He is also grateful to CIHR, NSERC, and SSHRC for supporting the tricouncil grant from the Canada-UK Artificial Intelligence (AI) Initiative “The self as agent-environment nexus: crossing disciplinary boundaries to help human selves and anticipate artificial selves” (ES/T01279X/1)(together with Karl J. Friston from the UK).

## Author Contributions

Y.Ç. and G.N. conceived the project. Y.Ç. analyzed the data, created figures and performed simulations. G.N. supervised the project. Y.Ç. and G.N. wrote the first draft. K.K., A.W., A.B. commented on the manuscript and contributed to revisions.

## Competing Interests

The authors declare no competing interests.

## SUPPLEMENTARY MATERIAL

### A Brief Introduction to Fluctuation-Dissipation Theorem

The fluctuation-dissipation theorem forms the basis of our hypothesis linking the autocorrelation function to event-related brain activity. While a full treatment is out of the scope of our paper, we will nonetheless provide a brief self-consistent introduction in our context. Interested reader can consult further references^1,2^.

We start by considering a generalized coordinate *x*(*t*) which can correspond to the firing rate of a cortical column. We denote the conjugate force to this coordinate as ℎ(*t*). To the leading order, the average response of 〈*x*(*t*)〉 to the force ℎ(*t*) is given by the Volterra expansion:

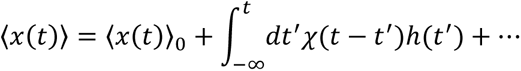

with 〈*x*(*t*)〉_0_ denoting the average value in the absence of any perturbation. If the system is highly nonlinear, then the dots in the expansion become non-negligible. However, to a leading order, *χ*(*t*) is enough to understand the response of the system to the external stimulus ℎ(*t*). The function *χ*(*t*) is called the linear response function. Our goal is to give an approximation for it.

Given the nature of event-related fields (ERFs) as average of neural activity across trials, we are interested in the expectation value 〈*x*(*t*)〉. Our first task is to describe a probability distribution that would give this average. We let *P*_0_[*x*(*t*)] denote the probability distribution of *x*(*t*) so that ∫ *dx*′(*t*)*P*_0_[*x*′(*t*)]*x*′(*t*) = 〈*x*(*t*)〉. We assume that there exists some constraint 〈*H*_0_[*x*(*t*)]〉 on the probability distribution. In statistical mechanics, this constraint can take the form of total energy^3^. In neuroscience, this function *H*_0_[*x*(*t*)] corresponds to empirical measurement of mean firing rates and covariances between neurons (see for example^4–6^, also^7^ for a more comprehensive overview). We will proceed by keeping *H*_0_[*x*(*t*)] general, without giving any specific information on what it is. Therefore our derivation will also be general across various choices for *H*_0_[*x*(*t*)]. Our only specification is that *H*_0_ is a function of *x*(*t*). The average 〈*H*_0_[*x*(*t*)]〉 will be given by the usual relation 〈*H*_0_[*x*(*t*)]〉 = ∫ *dx*^′(*t*)^*P*_0_[*x*′(*t*)]*H*_0_[*x*′(*t*)]. To obtain *P*_0_[*x*(*t*)], we will maximize the entropy *S* = −∫ *dx*^′(*t*)^*P*_0_[*x*′(*t*)] ln *P*_0_[*x*′(*t*)]. Using Lagrange multipliers, we can rewrite *S*′ = −*S* as

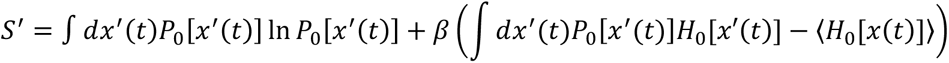

We can minimize *S*′ by taking its derivative with respect to *P*_0_[*x*(*t*)] and equate it to 0, which gives:

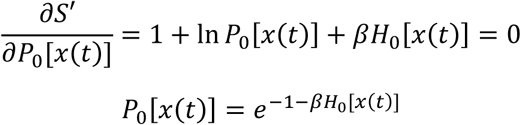

Note that we were careless about normalization of the probability distribution. We could have written one more Lagrange multiplier in the form of *α*(∫ *dx*′(*t*)*P*[*x*′(*t*)] − 1) which would add the normalization of the probability distribution to the derivation. Alternatively, we can construct a partition function *Z* = ∫ *dx*′(*t*)*e*^−*βH*_0_[*x*′(*t*)]^ and obtain the correct form of *P*_0_[*x*(*t*)] as

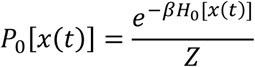

In the case of non-equilibrium phenomena such as event-related activity, we are interested in perturbations on *H*_0_. We will assume that our force ℎ(*t*) makes a change in *H*_0_. The neuroscientific intuition behind this assumption is the change of firing rates or correlation functions in various cognitive tasks in vivo or electrical stimulations in vitro. Let us denote the perturbed *H* as *H*[*x*(*t*), ℎ(*t*)]. For small ℎ(*t*), *H* can be approximated via a Taylor expansion around (*x*(*t*) = *x*(0), ℎ(*t*) = 0):

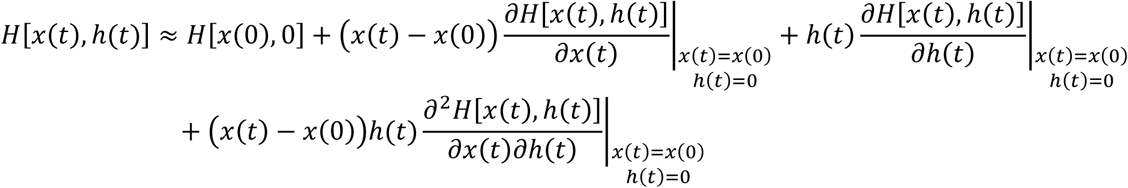

We will assume that under equilibrium conditions, *H*_0_ will be minimized. This corresponds to a saddle point approximation^3,8^. Since the probability distribution for *x*(*t*) is an exponential, the validity of approximation depends on how large *H*_0_ is. Nonetheless, it provides a good starting point. A further direction for this theory can be to evaluate the fluctuations around this saddle point and how they change the results. The advantage of this approximation is that it results in all the first derivatives being equal to 0 since if *H*_0_ is at an extremum, the first derivatives are equal to 0. Note that we are assuming that *H*[*x*(*t*), ℎ(*t*)] = *H*_0_[*x*(*t*)] + *δH*[*x*(*t*), ℎ(*t*)] where *δH*[*x*(0), 0] = 0 which gives us *H* = *H*_0_ in the absence of any external perturbation. Our final *H* is given by

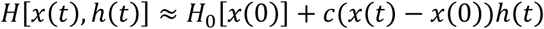

with the constant 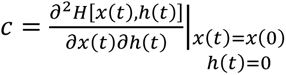. We note that in the formulation of statistical mechanics, *H* corresponds to energy and its changes are given by ∫ *dx*(*t*)*x*(*t*)ℎ(*t*) for generalized coordinates *x*(*t*) and conjugate forces ℎ(*t*). In this case, the saddle point approximation is not needed. We go with the explicit derivation to keep the theory more general.

External perturbation changes the probability distribution as well. We define the new probability distribution as *P*[*x*(*t*), ℎ(*t*)] = *W*[*x*(*t*)|*x*(0), ℎ(*t*)]*P*_0_[*x*(0), ℎ(*t*)] with transition probability *W*. The expectation value for *x*(*t*) is now given by

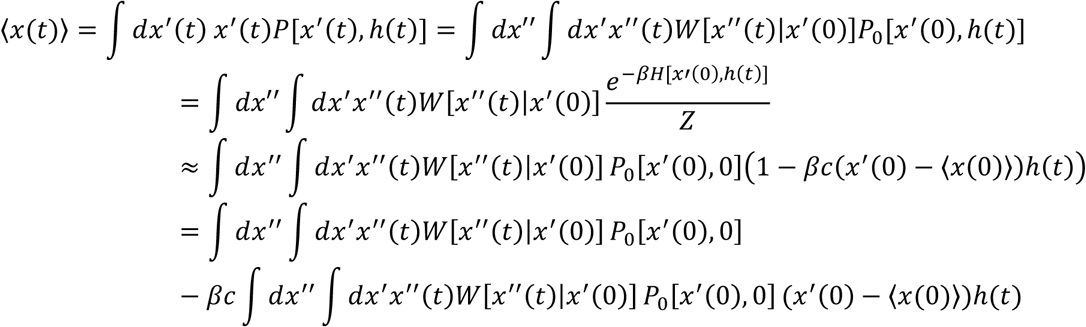

We can define the average values in the absence of perturbation as 〈*a*(*t*)〉_0_ = ∫ *dx*′′(*t*) ∫ *dx*′(*t*)*W*[*x*′(*t*)|*x*′′(0)]*P*_0_[*x*′′(0), 0]*a*(*t*):

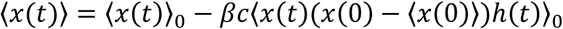

If we assume a constant force that is switched on at *t* = 0, ℎ(*t*) = −ℎ_0_*θ*(*t*), assume 〈*x*(0)〉 = 0 which corresponds to the common procedure of subtracting the mean of prestimulus period of an event-related potential and denote the autocorrelation function *A*(*τ*) = 〈*x*(*τ*)*x*(0)〉_0_ = 〈*x*(*t* − *τ*)*x*(*t*)〉_0_ (the last equality is valid under equilibrium conditions such as resting state with time translation invariance), we finally get

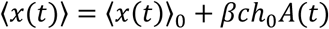

This interesting result gives an approximation of the task response purely from the fluctuations in the absence of any task. Going back to our initial problem of a linear response function, we identify

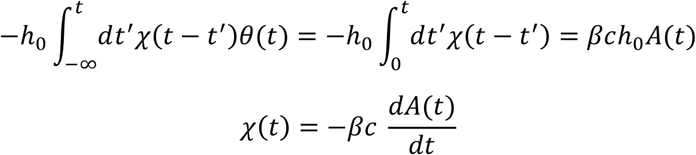

This formalism also enables us to put the intrinsic neural timescales on firm theoretical ground. Intrinsic neural timescales are defined as the time window during which prior information can influence the processing of new stimuli. It is clear that the autocorrelation function *A*(*t*) determines the time period in which the input can exert an influence on 〈*x*(*t*)〉. Since *A*(*t*) is usually of the exponential decay form and it is multiplicating the input ℎ, the time period in which *A*(*t*) is non-zero is also the time period in which different inputs can be added on top of each other, consistent with the definition of intrinsic neural timescales.

When the event-related potentials/fields are observed visually, obviously they do not correspond to the exact shape of the autocorrelation function. The divergence can be attributed to a number of potential reasons. Two potential reasons are mixing of sources from the brain and nonlinearity of the brain activity. Nonetheless, we can infer that the length of the intrinsic neural timescale should positively correlate with the magnitude of event-related response. This forms the basis of our hypothesis we investigate in the main manuscript.

Before we conclude this section, we note that there is one more way to show this relation through the usage of Martin-Siggia-Rose-De Dominicis-Janssen Path Integral^9–11^. This approach does not start with the maximum entropy assumption but rather uses a perturbation theoretical method on an arbitrarily general stochastic differential equation. Though the procedure is more abstract, the results are more general. We direct the interested reader to the references^12,13^.

### Additional Results from Spatiotemporal Cluster Testing

In this section, we present additional results for the spatiotemporal cluster permutation testing of event-related fields. Supplementary Table 1 shows the results of multiple comparisons of magnitude of event-related fields for identified clusters. Supplementary figures 1 to 4 show other clusters identified by spatiotemporal permutation testing.

**Supplementary Table 1.** Results of multiple comparisons of magnitude of event-related fields (mERFs)

| Cluster | mERF-1 | mERF-2 | Degrees of Freedom | t-statistic | Corrected p-value | Cohen's d |
| --- | --- | --- | --- | --- | --- | --- |
| Encode #1 | Happy | Sad | 110 | 0.311 | 0.756 | 0.059 |
| Encode #1 | Happy | Shape | 110 | 8.031 | <0.001 | 1.518 |
| Encode #1 | Sad | Shape | 110 | 7.087 | <0.001 | 1.339 |
| Encode #2 | Happy | Sad | 110 | -0.130 | 0.897 | -0.025 |
| Encode #2 | Happy | Shape | 110 | 8.295 | <0.001 | 1.568 |
| Encode #2 | Sad | Shape | 110 | 7.770 | <0.001 | 1.468 |
| Encode #3 | Happy | Sad | 110 | 0.343 | 0.732 | 0.065 |
| Encode #3 | Happy | Shape | 110 | 3.763 | 0.001 | 0.711 |
| Encode #3 | Sad | Shape | 110 | 3.549 | 0.001 | 0.671 |
| Encode #4 | Happy | Sad | 110 | -0.526 | 0.600 | -0.099 |
| Encode #4 | Happy | Shape | 110 | 4.500 | <0.001 | 0.850 |
| Encode #4 | Sad | Shape | 110 | 4.848 | <0.001 | 0.916 |
| Probe #1 | Happy | Sad | 110 | -0.193 | 0.847 | -0.036 |
| Probe #1 | Happy | Shape | 110 | 4.871 | <0.001 | 0.921 |
| Probe #1 | Sad | Shape | 110 | 5.029 | <0.001 | 0.950 |
| Probe #2 | Happy | Sad | 110 | 0.002 | 0.999 | 0.000 |
| Probe #2 | Happy | Shape | 110 | 5.309 | <0.001 | 1.003 |
| Probe #2 | Sad | Shape | 110 | 5.071 | <0.001 | 0.958 |

**Supplementary Figure 1.**
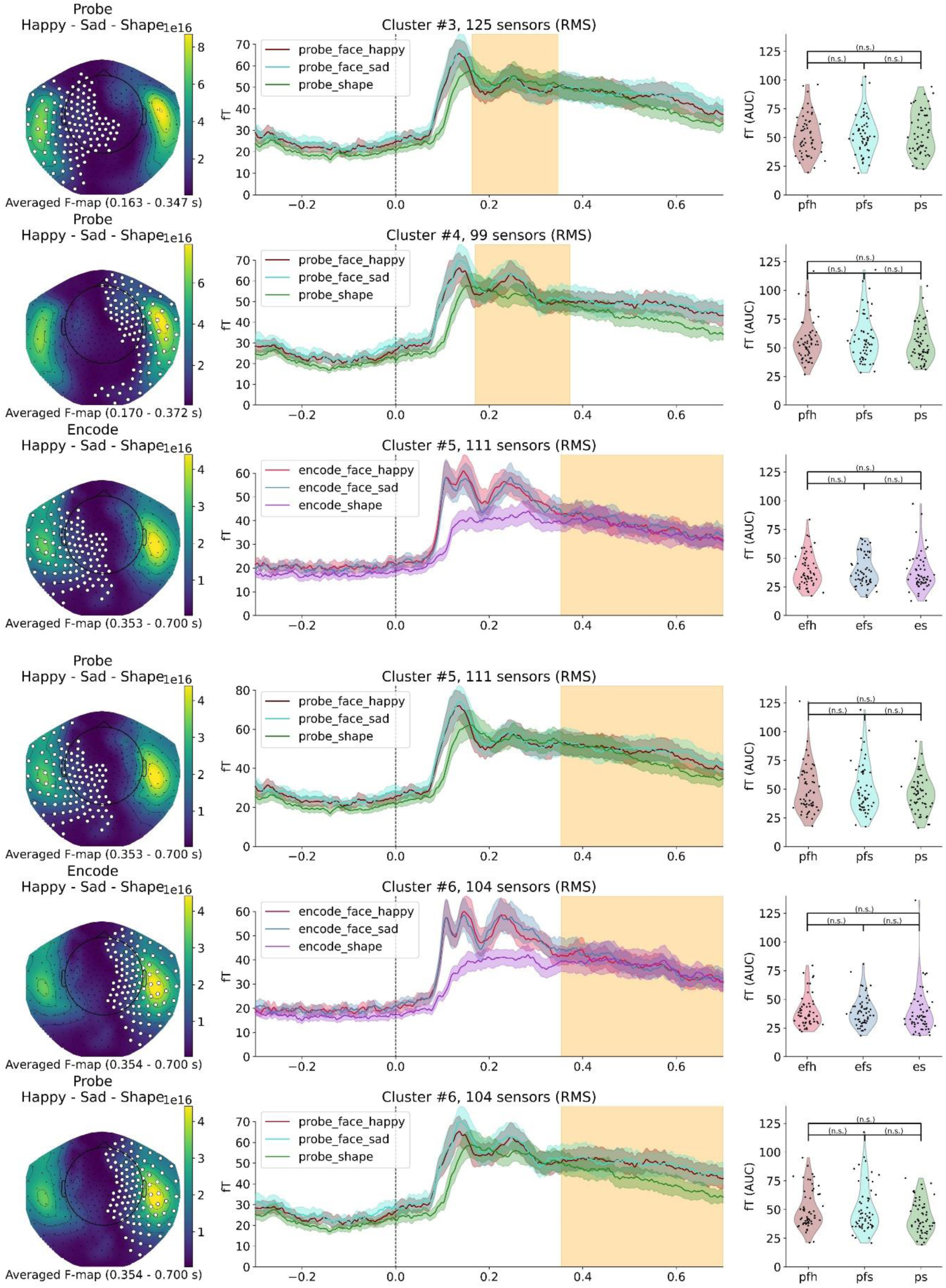
Non – Significant clusters from Happy versus Sad versus Shape Condition

**Supplementary Figure 2.**
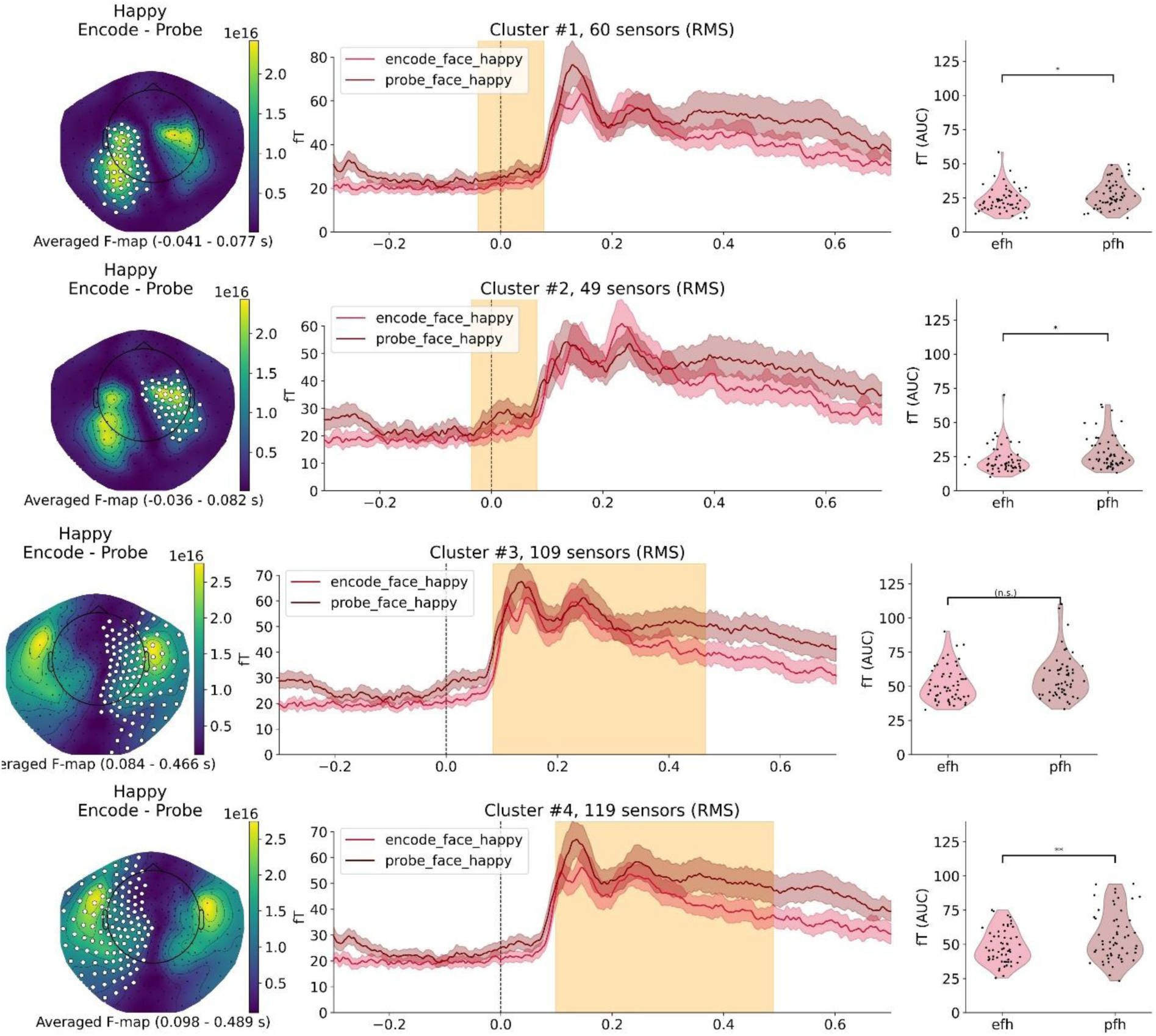
Encode versus Probe condition, in happy trials

**Supplementary Figure 3.**
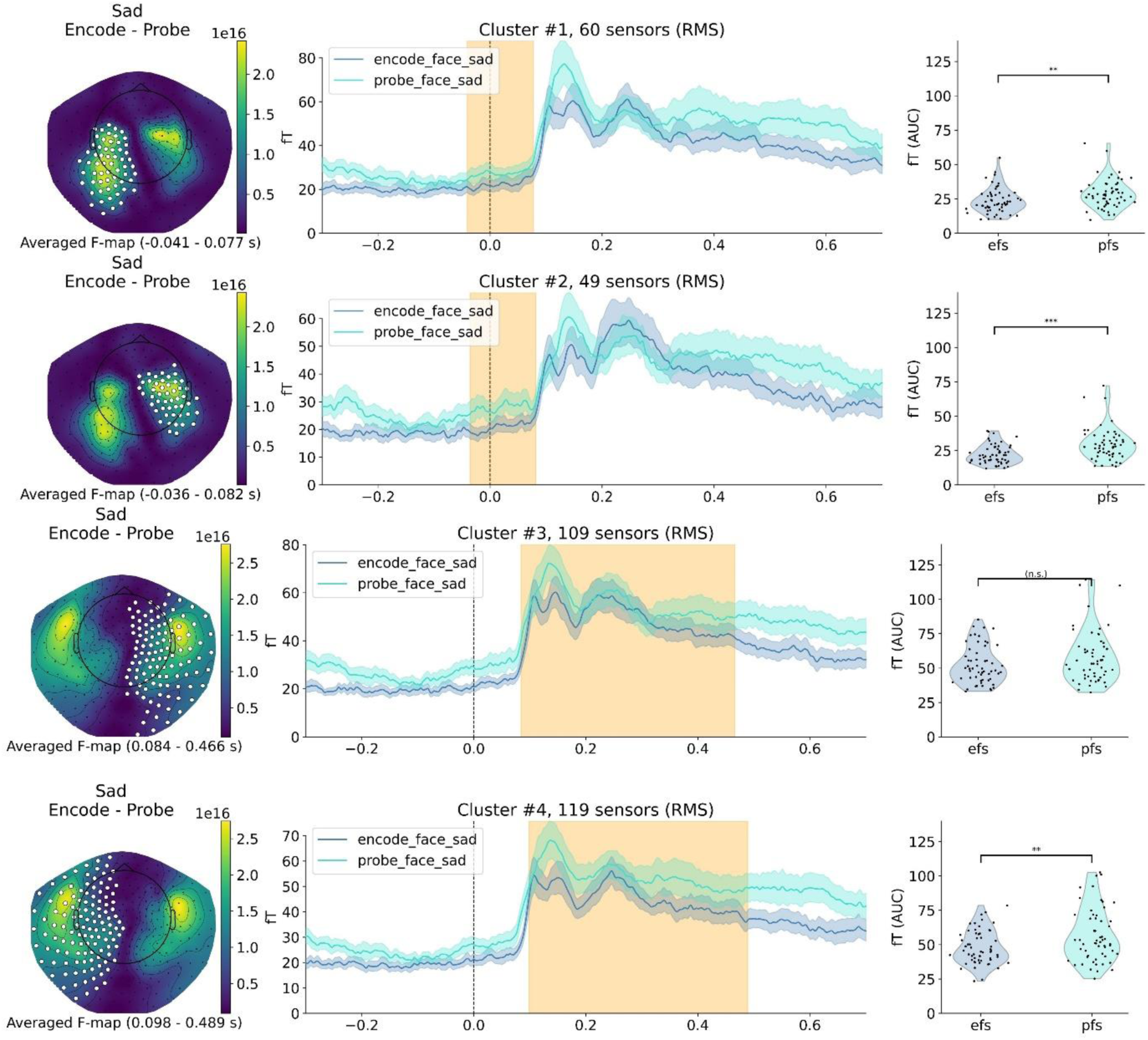
Encode versus Probe condition, in sad trials

**Supplementary Figure 4.**
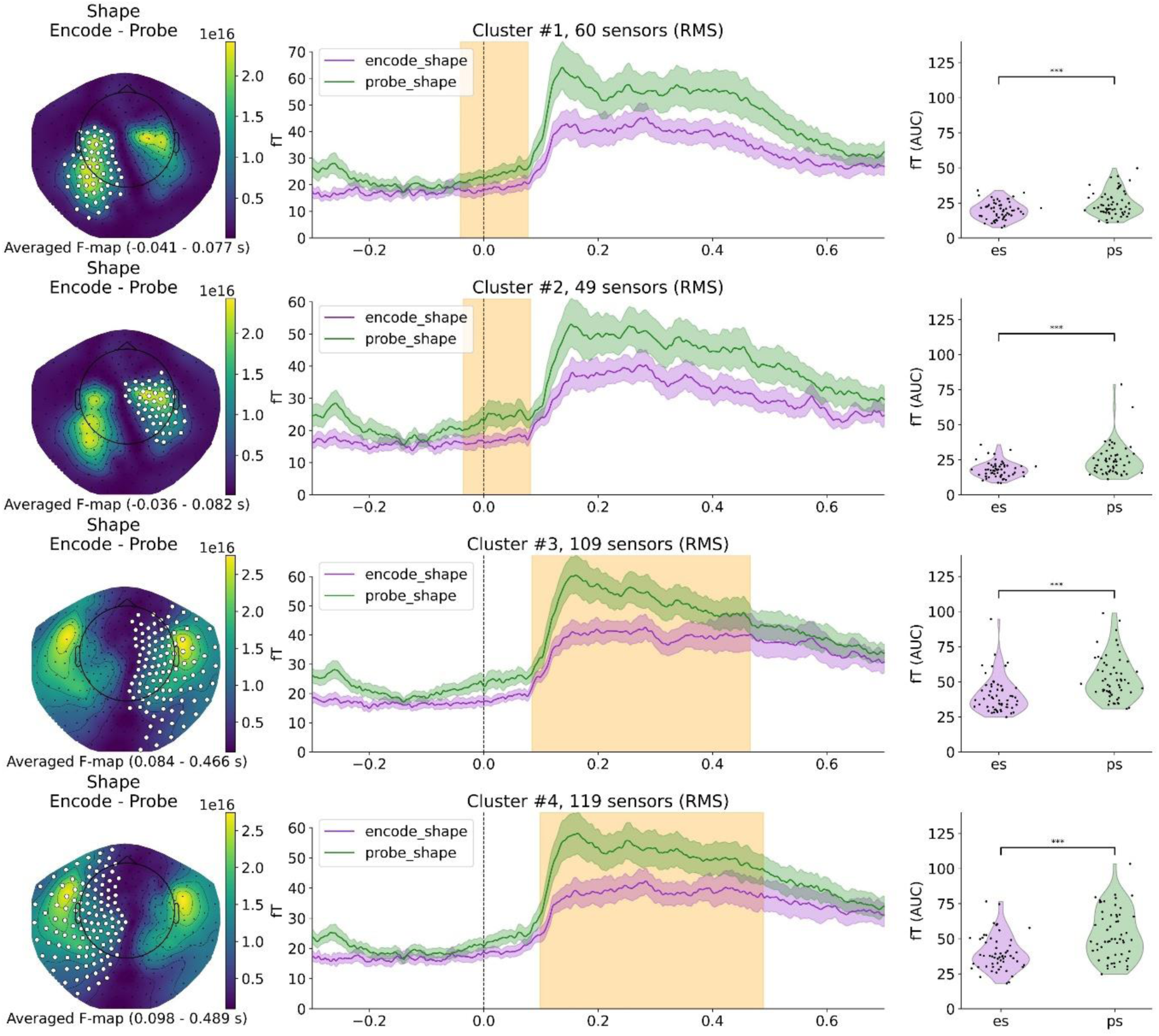
Encode versus Probe condition, in shape trials

### Forest Plots for the Relationship Between Reaction Times and Event-Related Fields

In addition to the relationship between the magnitude of event-related fields (mERFs) and intrinsic neural timescales (INTs), we investigated the relationship between these two measures with the behavioral outcome reaction times (RTs). Supplementary figure 5 shows the forest plot for the relationship between mERF and RT. We performed used a hierarchical Bayesian model with the same parameters for inference as in the main manuscript. The model specification is the same as

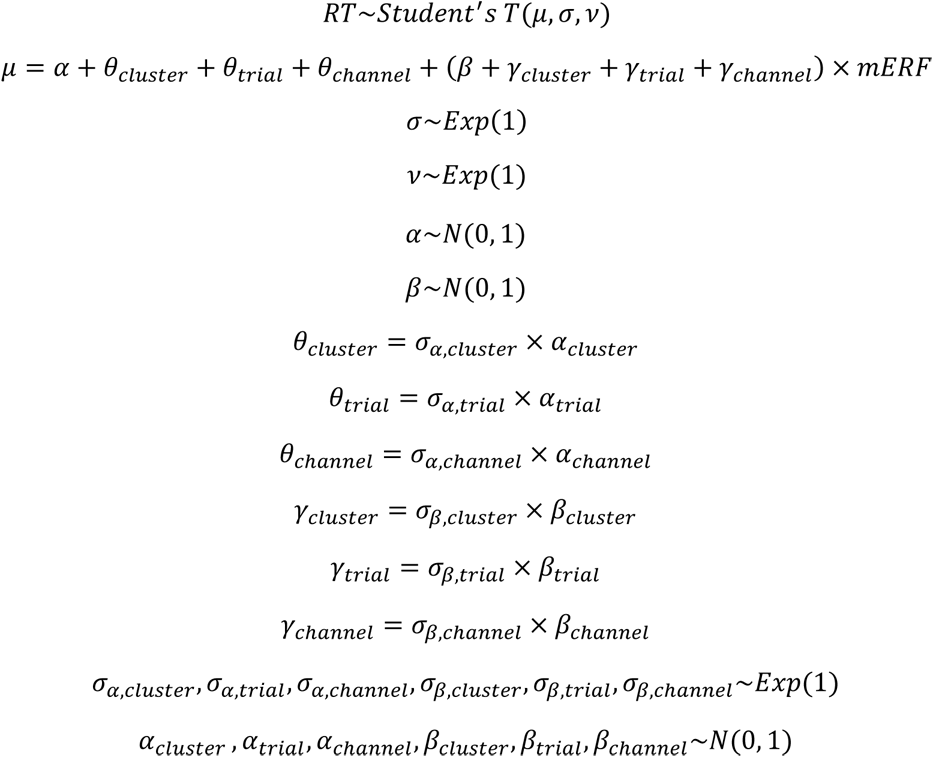

We observed a small negative effect for mERF (mean:-0.093, sd: 0.015, 94% HDI: [-0.124, −0.066], pd: 1, % in ROPE: 0.722). Supplementary figure 5a shows the forest plot for the fixed effect for mERF and random effects for clusters and trials. Supplementary figures 5b and 5c show the channel specific random effects and the sum of fixed effect and channel specific random effects respectively.

**Supplementary Figure 5.**
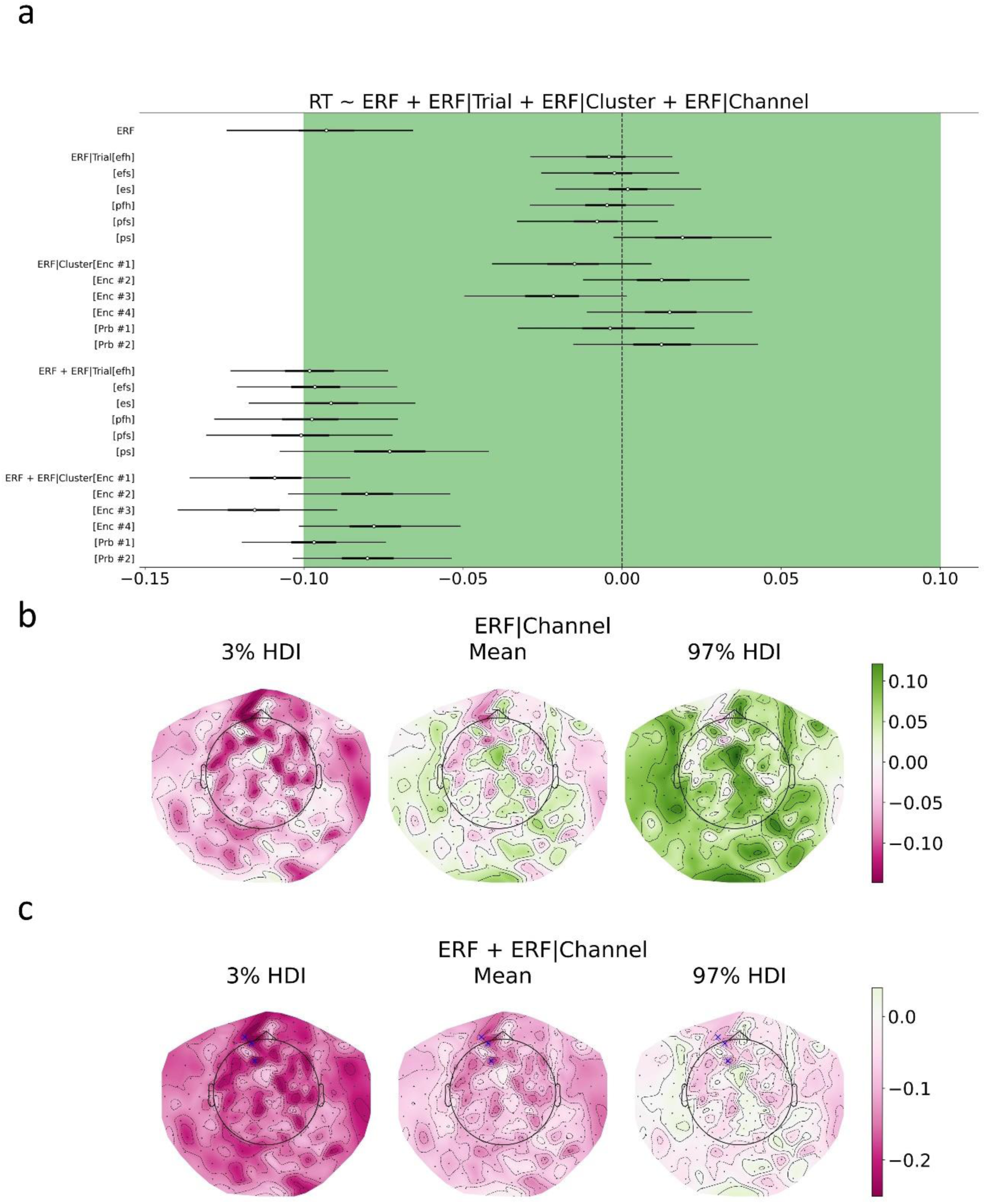
Relationship between reaction times and event-related fields. a. Coefficient estimates for a hierarchical Bayesian model where reaction time (RT) depends on magnitude of event-related fields (mERF) nested inside cluster, trial and channel variables. Dots denote the averages and lines denote 94% highest density probability intervals (HDI) of posterior distributions for coefficients. Limits for Region of Practical Equivalence (ROPE) is indicated with the green shading. b. Channel specific slopes estimated in the hierarchical model. c. Total effect for channels as a sum of fixed effect and random effect for each channel. In panels b and c, channels that have a ROPE less than 0.01 are indicated with a blue cross. Abbreviations: efh: encode face happy, efs: encode face sad, es: encode shape, pfh: probe face happy, pfs: probe face sad, ps: probe shape, Enc: Encode, Prb: Probe.

### Forest Plots for the Relationship Between Intrinsic Neural Timescales and Reaction Times

Finally, we analyzed the relationship between the autocorrelation windows (ACW) and reaction times (RT). Since ACWs are not nested in trial and cluster variables, we used a hierarchical model with one random effect: channels. The model specification is as follows:

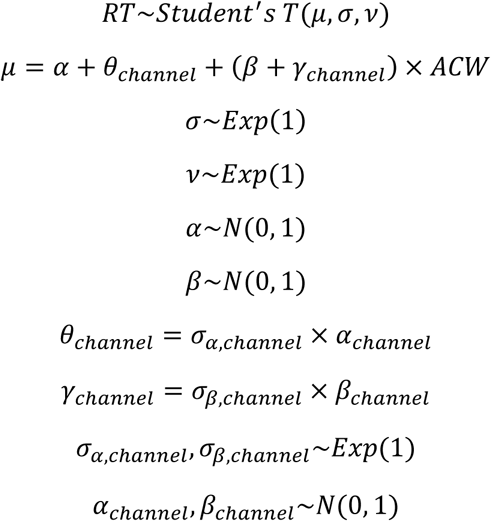

We observed a small and negative effect for ACW (mean: −0.105, sd: 0.004, 94% HDI: −0.112, −0.096, pd: 1, % in ROPE: 0.12). The same plot as above is shown below for fixed effect for ACW and random effects for channels:

---

**Supplementary Figure 6.**
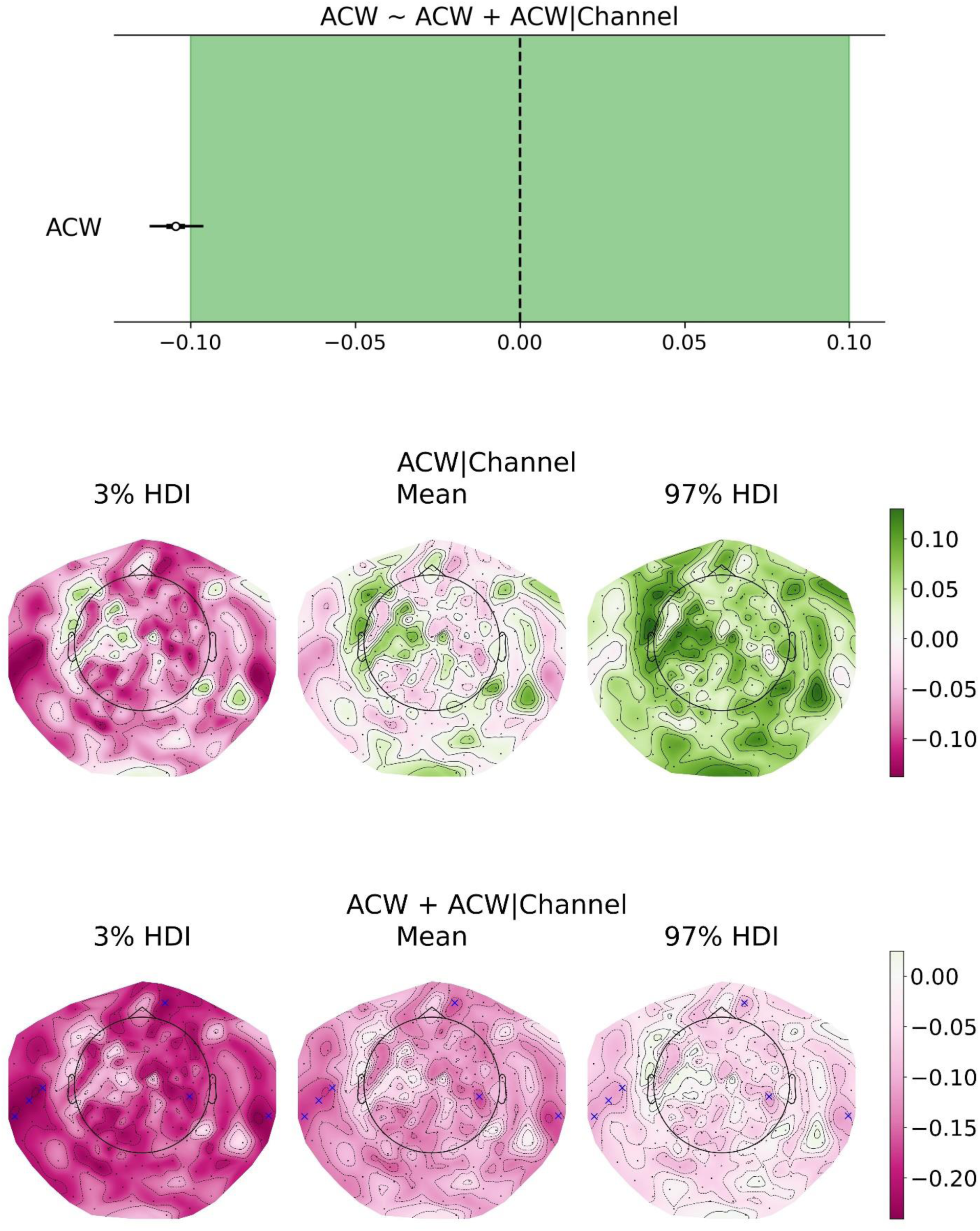
Relationship between reaction times and intrinsic neural timescales. a. Coefficient estimates for a hierarchical Bayesian model where reaction time (RT) depends on autocorrelation window (ACW) nested inside channel variable. Dots denote the averages and lines denote 94% highest density probability intervals (HDI) of posterior distributions for coefficients. Limits for Region of Practical Equivalence (ROPE) is indicated with the green shading. b. Channel specific slopes estimated in the hierarchical model. c. Total effect for channels as a sum of fixed effect and random effect for each channel. In panels b and c, channels that have a ROPE less than 0.01 are indicated with a blue cross. Abbreviations: efh: encode face happy, efs: encode face sad, es: encode shape, pfh: probe face happy, pfs: probe face sad, ps: probe shape, Enc: Encode, Prb: Probe.

## Notes

### Competing Interest Statement

The authors have declared no competing interest.

## References

1. Northoff, G. From Brain Dynamics to the Mind. (2024). Academic Press, Elsevier, Amsterdam, New York

2. Wolff, A. et al. Intrinsic neural timescales: temporal integration and segregation. Trends in Cognitive Sciences 26, 159–173 (2022).

3. Ito, T., Hearne, L. J. & Cole, M. W. A cortical hierarchy of localized and distributed processes revealed via dissociation of task activations, connectivity changes, and intrinsic timescales. NeuroImage 221, 117141 (2020).

4. Golesorkhi, M. et al. The brain and its time: intrinsic neural timescales are key for input processing. Commun Biol 4, 1–16 (2021).

5. Yeshurun, Y., Nguyen, M. & Hasson, U. The default mode network: where the idiosyncratic self meets the shared social world. Nat Rev Neurosci 22, 181–192 (2021).

6. Hasson, U., Chen, J. & Honey, C. J. Hierarchical process memory: memory as an integral component of information processing. Trends in Cognitive Sciences 19, 304–313 (2015).

7. Hasson, U., Yang, E., Vallines, I., Heeger, D. J. & Rubin, N. A Hierarchy of Temporal Receptive Windows in Human Cortex. J. Neurosci. 28, 2539–2550 (2008).

8. Murray, J. D. et al. A hierarchy of intrinsic timescales across primate cortex. Nat Neurosci 17, 1661–1663 (2014).

9. Wolff, A. et al. Neural variability quenching during decision-making: Neural individuality and its prestimulus complexity. NeuroImage 192, 1–14 (2019).

10. Smith, D., Wolff, A., Wolman, A., Ignaszewski, J. & Northoff, G. Temporal continuity of self: Long autocorrelation windows mediate self-specificity. NeuroImage 257, 119305 (2022).

11. Chang, C. H. C., Nastase, S. A. & Hasson, U. Information flow across the cortical timescale hierarchy during narrative construction. Proc. Natl. Acad. Sci. U.S.A. 119, e2209307119 (2022).

12. Zilio, F. et al. Altered brain dynamics index levels of arousal in complete locked-in syndrome. Commun Biol 6, 1–19 (2023).

13. Zilio, F. et al. Are intrinsic neural timescales related to sensory processing? Evidence from abnormal behavioral states. NeuroImage 226, 117579 (2021).

14. Buccellato, A. et al. Disrupted relationship between intrinsic neural timescales and alpha peak frequency during unconscious states - A high-density EEG study. Neuroimage 265, 119802 (2023).

15. Assecondi, S., Villa-Sánchez, B. & Shapiro, K. Event-related potentials as markers of efficacy for combined working memory training and transcranial direct current stimulation regimens: A proof-of-concept study. Frontiers in Systems Neuroscience 16, (2022).

16. O’Reilly, J. A., Wehrman, J. & Sowman, P. F. A Guided Tutorial on Modelling Human Event-Related Potentials with Recurrent Neural Networks. Sensors 22, 9243 (2022).

17. Shah, A. S. et al. Neural Dynamics and the Fundamental Mechanisms of Event-related Brain Potentials. Cerebral Cortex 14, 476–483 (2004).

18. Kolossa, A., Kopp, B. & Fingscheidt, T. A computational analysis of the neural bases of Bayesian inference. NeuroImage 106, 222–237 (2015).

19. Zeraati, R. et al. Intrinsic timescales in the visual cortex change with selective attention and reflect spatial connectivity. Nat Commun 14, 1858 (2023).

20. Fox, M. D. et al. The human brain is intrinsically organized into dynamic, anticorrelated functional networks. Proceedings of the National Academy of Sciences 102, 9673–9678 (2005).

21. Boly, M. et al. Baseline brain activity fluctuations predict somatosensory perception in humans. Proceedings of the National Academy of Sciences 104, 12187–12192 (2007).

22. Hesselmann, G., Kell, C. A., Eger, E. & Kleinschmidt, A. Spontaneous local variations in ongoing neural activity bias perceptual decisions. Proc Natl Acad Sci U S A 105, 10984–10989 (2008).

23. Mennes, M. et al. Inter-individual differences in resting-state functional connectivity predict task-induced BOLD activity. NeuroImage 50, 1690–1701 (2010).

24. Coste, C. P., Sadaghiani, S., Friston, K. J. & Kleinschmidt, A. Ongoing Brain Activity Fluctuations Directly Account for Intertrial and Indirectly for Intersubject Variability in Stroop Task Performance. Cerebral Cortex 21, 2612–2619 (2011).

25. He, B. J. Spontaneous and Task-Evoked Brain Activity Negatively Interact. J. Neurosci. 33, 4672–4682 (2013).

26. Sadaghiani, S. & Kleinschmidt, A. Functional interactions between intrinsic brain activity and behavior. NeuroImage 80, 379–386 (2013).

27. Sadaghiani, S., Poline, J.-B., Kleinschmidt, A. & D’Esposito, M. Ongoing dynamics in large-scale functional connectivity predict perception. Proceedings of the National Academy of Sciences 112, 8463–8468 (2015).

28. Sadaghiani, S., Hesselmann, G. & Kleinschmidt, A. Distributed and Antagonistic Contributions of Ongoing Activity Fluctuations to Auditory Stimulus Detection. J. Neurosci. 29, 13410–13417 (2009).

29. Huang, Z. et al. Is There a Nonadditive Interaction Between Spontaneous and Evoked Activity? Phase-Dependence and Its Relation to the Temporal Structure of Scale-Free Brain Activity. Cerebral Cortex 27, 1037–1059 (2017).

30. Çatal, Y., Gomez-Pilar, J. & Northoff, G. Intrinsic dynamics and topography of sensory input systems. Cerebral Cortex 32, 4592–4604 (2022).

31. Kamp, T., Sorger, B., Benjamins, C., Hausfeld, L. & Goebel, R. The prestimulus default mode network state predicts cognitive task performance levels on a mental rotation task. Brain and Behavior 8, e01034 (2018).

32. Wu, Y.-H., Podvalny, E., Levinson, M. & He, B. J. Network mechanisms of ongoing brain activity’s influence on conscious visual perception. Nat Commun 15, 5720 (2024).

33. Wolff, A. et al. Prestimulus dynamics blend with the stimulus in neural variability quenching. NeuroImage 238, 118160 (2021).

34. Kolvoort, I. R., Wainio-Theberge, S., Wolff, A. & Northoff, G. Temporal integration as “common currency” of brain and self-scale-free activity in resting-state EEG correlates with temporal delay effects on self-relatedness. Human Brain Mapping 41, 4355–4374 (2020).

35. Wainio-Theberge, S., Wolff, A. & Northoff, G. Dynamic relationships between spontaneous and evoked electrophysiological activity. Commun Biol 4, 1–17 (2021).

36. Waschke, L., Wöstmann, M. & Obleser, J. States and traits of neural irregularity in the age-varying human brain. Sci Rep 7, 17381 (2017).

37. Iemi, L., Chaumon, M., Crouzet, S. M. & Busch, N. A. Spontaneous Neural Oscillations Bias Perception by Modulating Baseline Excitability. J Neurosci 37, 807–819 (2017).

38. Iemi, L. et al. Ongoing neural oscillations influence behavior and sensory representations by suppressing neuronal excitability. NeuroImage 247, 118746 (2022).

39. Romei, V. et al. Spontaneous fluctuations in posterior alpha-band EEG activity reflect variability in excitability of human visual areas. Cereb Cortex 18, 2010–2018 (2008).

40. Northoff, G., Zilio, F. & Zhang, J. Beyond task response-Pre-stimulus activity modulates contents of consciousness. Phys Life Rev 49, 19–37 (2024).

41. Braun, W., Matsuzaka, Y., Mushiake, H., Northoff, G. & Longtin, A. Non-additive activity modulation during a decision making task involving tactic selection. Cogn Neurodyn 16, 117–133 (2022).

42. Churchland, M. M. et al. Stimulus onset quenches neural variability: a widespread cortical phenomenon. Nat Neurosci 13, 369–378 (2010).

43. Petersen, C. C. H., Hahn, T. T. G., Mehta, M., Grinvald, A. & Sakmann, B. Interaction of sensory responses with spontaneous depolarization in layer 2/3 barrel cortex. Proceedings of the National Academy of Sciences 100, 13638–13643 (2003).

44. Balakrishnan, V. Elements of Nonequilibrium Statistical Mechanics. (Springer International Publishing, Cham, 2021). doi:10.1007/978-3-030-62233-6.

45. Callen, H. B. & Welton, T. A. Irreversibility and Generalized Noise. Phys. Rev. 83, 34– 40 (1951).

46. David Tong: Kinetic Theory. https://www.damtp.cam.ac.uk/user/tong/kinetic.html.

47. Jansen, B. H. & Rit, V. G. Electroencephalogram and visual evoked potential generation in a mathematical model of coupled cortical columns. Biol. Cybern. 73, 357–366 (1995).

48. Jansen, B. H., Zouridakis, G. & Brandt, M. E. A neurophysiologically-based mathematical model of flash visual evoked potentials. Biol Cybern 68, 275–283 (1993).

49. David, O. & Friston, K. J. A neural mass model for MEG/EEG: coupling and neuronal dynamics. Neuroimage 20, 1743–1755 (2003).

50. David, O., Harrison, L. & Friston, K. J. Modelling event-related responses in the brain. Neuroimage 25, 756–770 (2005).

51. Kiebel, S. J., David, O. & Friston, K. J. Dynamic causal modelling of evoked responses in EEG/MEG with lead field parameterization. Neuroimage 30, 1273–1284 (2006).

52. Wang, X.-J. Macroscopic gradients of synaptic excitation and inhibition in the neocortex. Nat Rev Neurosci 21, 169–178 (2020).

53. Nugent, A. C. et al. The NIMH intramural healthy volunteer dataset: A comprehensive MEG, MRI, and behavioral resource. Sci Data 9, 518 (2022).

54. Herrmann, L. et al. fMRI Revealed Reduced Amygdala Activation after Nx4 in Mildly to Moderately Stressed Healthy Volunteers in a Randomized, Placebo-Controlled, Cross-Over Trial. Sci Rep 10, 3802 (2020).

55. Cornwell, B. R. et al. Evoked amygdala responses to negative faces revealed by adaptive MEG beamformers. Brain Res 1244, 103–112 (2008).

56. Varkevisser, T. et al. Pattern classification based on the amygdala does not predict an individual’s response to emotional stimuli. Hum Brain Mapp 44, 4452–4466 (2023).

57. DeFelipe, J., Alonso-Nanclares, L. & Arellano, J. I. Microstructure of the neocortex: comparative aspects. J Neurocytol 31, 299–316 (2002).

58. Miller, K. D. Understanding layer 4 of the cortical circuit: a model based on cat V1. Cereb Cortex 13, 73–82 (2003).

59. Rackauckas, C. & Nie, Q. Adaptive methods for stochastic differential equations via natural embeddings and rejection sampling with memory. DCDS-B 22, 2731–2761 (2017).

60. Rackauckas, C. & Nie, Q. DifferentialEquations.jl – A Performant and Feature-Rich Ecosystem for Solving Differential Equations in Julia. Journal of Open Research Software 5, (2017).

61. Rackauckas, C. & Nie, Q. Stability-Optimized High Order Methods and Stiffness Detection for Pathwise Stiff Stochastic Differential Equations. Preprint at 10.48550/arXiv.1804.04344 (2018).

62. Kennedy, C. A. & Carpenter, M. H. Additive Runge–Kutta schemes for convection– diffusion–reaction equations. Applied Numerical Mathematics 44, 139–181 (2003).

63. Khintchine, A. Korrelationstheorie der stationären stochastischen Prozesse. Math. Ann. 109, 604–615 (1934).

64. Jas, M., Engemann, D., Raimondo, F., Bekhti, Y. & Gramfort, A. Automated rejection and repair of bad trials in MEG/EEG. in 6th International Workshop on Pattern Recognition in Neuroimaging (PRNI) (Trento, Italy, 2016).

65. Jas, M., Engemann, D. A., Bekhti, Y., Raimondo, F. & Gramfort, A. Autoreject: Automated artifact rejection for MEG and EEG data. NeuroImage 159, 417–429 (2017).

66. Hämäläinen, M. S. & Ilmoniemi, R. J. Interpreting magnetic fields of the brain: minimum norm estimates. Med Biol Eng Comput 32, 35–42 (1994).

67. Veillette, J. P., Heald, S. L. M., Wittenbrink, B., Reis, K. S. & Nusbaum, H. C. Single-trial visually evoked potentials predict both individual choice and market outcomes. Sci Rep 13, 14340 (2023).

68. van Bijnen, S., Muotka, J. & Parviainen, T. Divergent auditory activation in relation to inhibition task performance in children and adults. Hum Brain Mapp 44, 4972–4985 (2023).

69. Foerster, F. R., Chidharom, M. & Giersch, A. Enhanced temporal resolution of vision in action video game players. NeuroImage 269, 119906 (2023).

70. Gramfort, A. et al. MEG and EEG data analysis with MNE-Python. Front. Neurosci. 7, (2013).

71. Delorme, A., Sejnowski, T. & Makeig, S. Enhanced detection of artifacts in EEG data using higher-order statistics and independent component analysis. Neuroimage 34, 1443–1449 (2007).

72. Abril-Pla, O. et al. PyMC: a modern, and comprehensive probabilistic programming framework in Python. PeerJ Comput Sci 9, e1516 (2023).

73. Salvatier, J., Wiecki, T. & Fonnesbeck, C. Probabilistic Programming in Python using PyMC. Preprint at 10.48550/arXiv.1507.08050 (2015).

74. Kumar, R., Carroll, C., Hartikainen, A. & Martin, O. ArviZ a unified library for exploratory analysis of Bayesian models in Python. Journal of Open Source Software 4, 1143 (2019).

75. Capretto, T., et al. Bambi: A simple interface for fitting Bayesian linear models in Python. Preprint at 10.48550/arXiv.2012.10754 (2022).

76. Hoffman, M. D. & Gelman, A. The No-U-Turn Sampler: Adaptively Setting Path Lengths in Hamiltonian Monte Carlo. Preprint at 10.48550/arXiv.1111.4246 (2011).

77. Gelman, A. & Rubin, D. B. Inference from Iterative Simulation Using Multiple Sequences. Statistical Science 7, 457–472 (1992).

78. Çatal, Y. Intracolumnar Excitatory and Inhibitory Connections Relate Intrinsic Neural Timescales to Task Related Activity.

79. Makowski, D., Ben-Shachar, M. S., Chen, S. H. A. & Lüdecke, D. Indices of Effect Existence and Significance in the Bayesian Framework. Front. Psychol. 10, (2019).

80. Kruschke, J. K. Rejecting or accepting parameter values in Bayesian estimation. Advances in Methods and Practices in Psychological Science 1, 270–280 (2018).

81. Felleman, D. J. & Van Essen, D. C. Distributed hierarchical processing in the primate cerebral cortex. Cereb Cortex 1, 1–47 (1991).

82. Honey, C. J. et al. Slow Cortical Dynamics and the Accumulation of Information over Long Timescales. Neuron 76, 423–434 (2012).

83. Hariri, A. R., Mattay, V. S., Tessitore, A., Fera, F. & Weinberger, D. R. Neocortical modulation of the amygdala response to fearful stimuli. Biol Psychiatry 53, 494–501 (2003).

84. McElreath, R. Statistical Rethinking: A Bayesian Course with Examples in R and STAN. (Chapman and Hall/CRC, Boca Raton London New York, 2020).

85. Wolman, A. et al. Intrinsic neural timescales mediate the cognitive bias of self – temporal integration as key mechanism. NeuroImage 268, 119896 (2023).

86. Gao, R., van den Brink, R. L., Pfeffer, T. & Voytek, B. Neuronal timescales are functionally dynamic and shaped by cortical microarchitecture. eLife 9, e61277 (2020).

87. Raut, R. V., Snyder, A. Z. & Raichle, M. E. Hierarchical dynamics as a macroscopic organizing principle of the human brain. Proceedings of the National Academy of Sciences 117, 20890–20897 (2020).

88. Faisal, A. A., Selen, L. P. J. & Wolpert, D. M. Noise in the nervous system. Nat Rev Neurosci 9, 292–303 (2008).

89. Schneidman, E., Freedman, B. & Segev, I. Ion channel stochasticity may be critical in determining the reliability and precision of spike timing. Neural Comput 10, 1679–1703 (1998).

90. White, J. A. et al. Channel noise in neurons. Trends in Neurosciences 23, 131–137 (2000).

91. Hernan, M. A. & Robins, J. M. Causal Inference: What If. (CRC Press, 2024).

## REFERENCES

1. Balakrishnan, V. Elements of Nonequilibrium Statistical Mechanics. (Springer International Publishing, Cham, 2021). doi:10.1007/978-3-030-62233-6.

2. David Tong: Kinetic Theory. https://www.damtp.cam.ac.uk/user/tong/kinetic.html.

3. Kardar, M. Statistical Physics of Particles. (Cambridge University Press, 2007).

4. Schneidman, E., Berry, M. J., Segev, R. & Bialek, W. Weak pairwise correlations imply strongly correlated network states in a neural population. Nature 440, 1007–1012 (2006).

5. Tkačik, G. et al. Searching for Collective Behavior in a Large Network of Sensory Neurons. PLOS Computational Biology 10, e1003408 (2014).

6. Meshulam, L., Gauthier, J. L., Brody, C. D., Tank, D. W. & Bialek, W. Collective Behavior of Place and Non-place Neurons in the Hippocampal Network. Neuron 96, 1178–1191.e4 (2017).

7. Bialek, W. Biophysics: Searching for Principles. (Princeton University Press, 2012).

8. Bender, C. M. & Orszag, S. A. Advanced Mathematical Methods for Scientists and Engineers I: Asymptotic Methods and Perturbation Theory. (Springer Science & Business Media, 1999).

9. Martin, P. C., Siggia, E. D. & Rose, H. A. Statistical Dynamics of Classical Systems. Phys. Rev. A 8, 423–437 (1973).

10. De Dominicis, C. & Peliti, L. Field-theory renormalization and critical dynamics above ${T}_{c}$: Helium, antiferromagnets, and liquid-gas systems. Phys. Rev. B 18, 353–376 (1978).

11. De Dominicis, C. Technics of field renormalization and dynamics of critical phenomena. J Phys (Paris), Colloq, (1), C1247-C1253. (1976).

12. Chow, C. C. & Buice, M. A. Path Integral Methods for Stochastic Differential Equations. The Journal of Mathematical Neuroscience (JMN) 5, 8 (2015).

13. Helias, M. & Dahmen, D. Statistical Field Theory for Neural Networks. vol. 970 (2020).

